# Evolution of pH-sensitive transcription termination during adaptation to repeated long-term starvation

**DOI:** 10.1101/2024.03.01.582989

**Authors:** Sarah B. Worthan, Robert D. P. McCarthy, Mildred Delaleau, Ryan Stikeleather, Benjamin P. Bratton, Marc Boudvillain, Megan G. Behringer

## Abstract

Fluctuating environments that consist of regular cycles of co-occurring stress are a common challenge faced by cellular populations. For a population to thrive in constantly changing conditions, an ability to coordinate a rapid cellular response is essential. Here, we identify a mutation conferring an arginine-to-histidine (Arg to His) substitution in the transcription terminator Rho. The *rho* R109H mutation frequently arose in *E. coli* populations experimentally evolved under repeated long-term starvation conditions, during which feast and famine result in drastic environmental pH fluctuations. Metagenomic sequencing revealed that populations containing the *rho* mutation also possess putative loss-of-function mutations in *ydcI*, which encodes a recently characterized transcription factor associated with pH homeostasis. Genetic reconstructions of these mutations show that the *rho* allele confers a plastic alkaline-induced reduction of Rho function that, when found in tandem with a Δ*ydcI* allele, leads to intracellular alkalinization and genetic assimilation of Rho mutant function. We further identify Arg to His substitutions at analogous sites in *rho* alleles from species originating from fluctuating alkaline environments. Our results suggest that Arg to His substitutions in global regulators of gene expression can serve to rapidly coordinate complex responses through pH sensing and shed light on how cellular populations across the tree of life use environmental cues to coordinate rapid responses to complex, fluctuating environments.

## Introduction

Cellular populations regularly encounter fluctuating environmental stressors in their natural contexts, such as changes in resource availability, temperature, pH, and oxidative stress ^1–3^. This holds particularly true for microorganisms, which have evolved to thrive in some of Earth’s most extreme and fluctuation prone environments. Even in the most forgiving of habitats, microbial growth is primarily limited due, in part, to the scarcity of carbon and nitrogen sources and competition between microbes ^4–6^. Given that resource availability fluctuates temporally and spatially in natural environments ^3^, microbial growth in nature occurs in bursts as resources become accessible, resulting in long, stagnant periods of starvation punctuated by transient periods of growth, commonly referred to as feast/famine cycles ^4,7^. Previous work has revealed several common microbial responses to persistent famine conditions, particularly those experienced by batch cultures during long-term stationary phase. These responses include an increase in genetic diversity, elevated mutation rates, morphological changes, and the emergence of the <u>g</u>rowth <u>a</u>dvantage in <u>s</u>tationary <u>p</u>hase (GASP) phenotype which confers an increased ability to catabolize alternative energy sources, such as amino acids, present in necromass as well as modulation of the stress response ^8–14^. Building upon this, recent work suggests that bacteria exhibit distinct responses depending on the frequency and magnitude of nutrient resource fluctuations, indicating that these fluctuations are an important determinant of bacterial behavior ^3,15–17^. For instance, the duration of starvation between resource replenishment can influence evolutionary outcomes; a narrow interval between resource replenishment events leads to the emergence of ecotypes within populations, while populations evolving in fluctuating environments with greater durations between resource replenishment events experience rapid selective sweeps and exhibit a high level of mutational order ^16,18^. Notably, microbial populations subjected to cycles of prolonged starvation followed by a rapid influx of resources (i.e. 100 day feast/famine cycles) still exhibit GASP-like behavior but evolve mutations that are distinct from the mutations traditionally associated with the GASP phenotype ^18^. Thus, further investigation of how bacteria adapt to repeated nutrient resource fluctuations is critical to contextualizing microbial behavior in the wild.

Among the complications that microbes face in the wild, the reintroduction of nutrient resources into environments presents an inherent complex stress. Here, microbes must contend with additional stressors that coincide with nutrient resource replenishment, such as shifts in pH. Microbes are further challenged as they metabolize the newly introduced resources since excreted metabolic byproducts can result in further environmental changes. These ecological feedbacks are also common in laboratory environments, particularly the significant pH fluctuations that can accompany growth ^19^. For example, when *E. coli* populations are cultivated in the amino acid-based LB broth, their metabolic activities induce remarkable shifts in pH ^20,21^. As these bacteria metabolize the amino acids for carbon, they release ammonia, causing the medium pH to undergo a dramatic transformation, shifting from a neutral pH of 7.0 to an alkaline pH of 9.0 in approximately 24 hours ^22,23^.

Consequently, many bacterial species have evolved mechanisms to sense and rapidly respond to changes in proton concentrations ^24^. For pathogens and marine bacteria which are frequently exposed to drastic shifts in external pH, the ability to sense these changes and respond through phenotypically plastic traits is critical to their survival ^25–28^. Often this ability to sense and respond to changes in pH are attributed to two-component regulatory systems, such as the MtrAB system in *Actinobacteria* ^29^ or the BatRS system in *Bartonella*. Another more universal plastic mechanism for pH sensing is the modulation of protein function via the protonation state of histidine residues, which is present across the Tree of Life. Histidine residues are often situated within functionally important sites of proteins, where their protonation status dictates the protein’s functionality ^30–32^. The protonation state of histidine residues also regulates the activity of particular cell-surface receptors and transporters, such as G-coupled receptors ^33^ and ion transporters ^34,35^. Moreover, tumorigenic arginine to histidine mutations are reported to contribute to cancers where they often confer pH sensing abilities ^36,37^. Thus, histidine residues, acting as pH sensors, can serve as a plastic mechanism to enable rapid responses to changing environments.

One powerful approach for investigating how organisms navigate complex stresses and identifying the resulting mutations contributing to adaptation is experimental evolution ^38,39^. Phenotypically plastic traits can evolve rapidly in evolution experiments and through these investigations, it has become evident that genes encoding global regulators of gene expression are frequent targets of selection when adapting to diverse environments, primarily due to their widespread impacts on various biological processes ^40,41^. One such essential global regulator is Rho, which terminates transcription at hundreds of sites throughout the *E. coli* genome ^42–45^. Rho-dependent termination plays critical cellular roles, including the delineation of transcription unit boundaries and surveillance of the gene expression machinery ^46^, as well as involvement in many conditional gene regulatory mechanisms ^47,48^. Additionally, Rho prevents unwanted transcription from loci antisense to genes or horizontally acquired genes, such as toxic prophage genes ^42,43^. Functioning as a ring-shaped homo-hexamer, Rho utilizes its crown-like primary binding site to bind nascent transcripts at poorly conserved C-rich regions called *rut* (<u>R</u>ho <u>ut</u>ilization) sites. Once bound to a *rut* site, the Rho hexamer guides the transcript into its central ring channel, where its secondary binding site mediates ATP-dependent [5’ to 3’] movement along the RNA chain until Rho catches up with and dissociates RNA polymerase, effectively terminating transcription ^49,50^.

Given the widespread prevalence of Rho-dependent termination in *E. coli*, any alteration in Rho function has the propensity to trigger extensive physiological changes. As such, numerous previous studies employing adaptive laboratory evolution under various conditions have reported nonsynonymous mutations in the *rho* gene. These conditions range from adaptation to heat stress ^51–54^ and exposure to copper ^55^ and ethanol ^56,57^ to the utilization of various carbon sources ^58,59^. Notably, previously reported adaptive mutations in *rho* exhibit variability, affecting residues distributed throughout the gene, particularly in regions associated with its core functions, including the primary and secondary binding site domains. Hence, the pleiotropic nature of Rho positions it as a frequently positively selected genetic target in *E. coli* when confronted with novel and intricate environmental challenges.

Here, we investigate a mutation in *rho* that results in an R109H substitution that fixed in 5 out of 16 *E. coli* populations during experimental evolution in repeated long-term starvation (RLTS) conditions consisting of 100 days between transfers into fresh LB-Miller broth ^18^. Each of these populations also contained putative loss of function mutations in an additional gene, *ydcI*, which encodes a largely uncharacterized transcriptional regulator related to pH homeostasis. Since arginine-to-histidine substitutions can alter a protein’s affinity to bind nucleic acids and are shown to grant pH-dependent activity to transcription factors in cancer ^37^, we hypothesize that the activity of Rho^R109H^ exhibits a similar pH-dependence that serves as a plastic adaptation to the neutral/alkaline fluctuations experienced during repeated starvation in LB broth. Using a combination of *in vitro* biochemical assays, fluorescence microscopy, and competitive co-culture, we examined how *rho* R109H and *ΔydcI* mutations contribute to the physiological adaptation of *E. coli* during repeated starvation. Additionally, through bioinformatic approaches, we further identify *rho* R109H alleles in wild bacterial strains that regularly experience neutral/alkaline fluctuations and infer amino acid residues of structural and functional importance within YdcI.

## Results

### Extensive parallel mutation in *rho* and *ydcI* during repeated long-term starvation

To gain insight into how microbes adapt to recurrent nutrient resource fluctuations following prolonged periods of famine, we evolved 16 *Escherichia coli* populations in parallel to a repeated long-term starvation (RLTS) regime of 100-day feast/famine cycles for a total of 900 days ^18^. Every 100 days, we transferred 1 mL of the aged culture into 9 mL of fresh LB-Miller broth, replenishing each culture’s resources. At this time, pretransfer aliquots of the populations were frozen to maintain a historical record of the evolution experiment. Growth in LB broth is supported through the catabolism of amino acids and leads to alkalinization of the medium within 24 hours which remains between pH 8 - pH 9 until additional resources are provided ^21,22,60^. As such, *E. coli* cultures are subjected to resource-depleted, alkaline conditions for the vast majority of the RLTS regime, essentially 99 days (**Fig S1A**). Every 100 days, when cultures are replenished with resources, the medium pH decreases dramatically from approximately pH 9 to pH 7, before slowly returning to an alkaline pH over the next 24 hours. Thus, populations adapted to these conditions must not only survive extended starvation under alkaline conditions, but also cope with the sudden downward shift in pH and repletion of resources concurrently.

Metagenomic sequencing throughout the 900-day experiment revealed that approximately half (7/16) of the populations evolved fixed mutations in *rho*, which encodes the Rho-dependent transcription termination factor (**Fig 1A; Fig S2A**). The majority of *rho* mutations (5/7) occurred in the first half of the RLTS experiment and swept to fixation within the first 400 days (**Fig 1B**). By 700 days, *rho* mutations were fixed in a total of 7 populations after which time no additional *rho* mutations were detected in the remaining 9 populations. All identified *rho* mutations occur within the primary RNA binding site of Rho. Thus, we suspect these mutations are likely to perturb interactions between Rho and its RNA target (**Fig 1C & 1E; Fig S3**). Furthermore, all 5 *rho* mutations that are fixed within the first 400 days result in an R109H nonsynonymous substitution, suggesting it may contribute to adaptation in RLTS conditions. After reconstructing the *rho* R109H allele into experimental ancestor background, we found that the R109H allele did not confer any considerable growth effects in conditions that previously produced other experimentally evolved *rho* mutations, including heat stress ^52^, exposure to ethanol ^57^, and osmotic stress ^61^ (**Fig S4**). This lack of phenotype is consistent with Rho’s role as a pleiotropic “master-regulator” ^62^ and indicates that any phenotypes associated with the *rho* R109H allele may be contingent on specific conditions that occur during RTLS or dependent on an earlier arising mutation.

**Fig 1:**
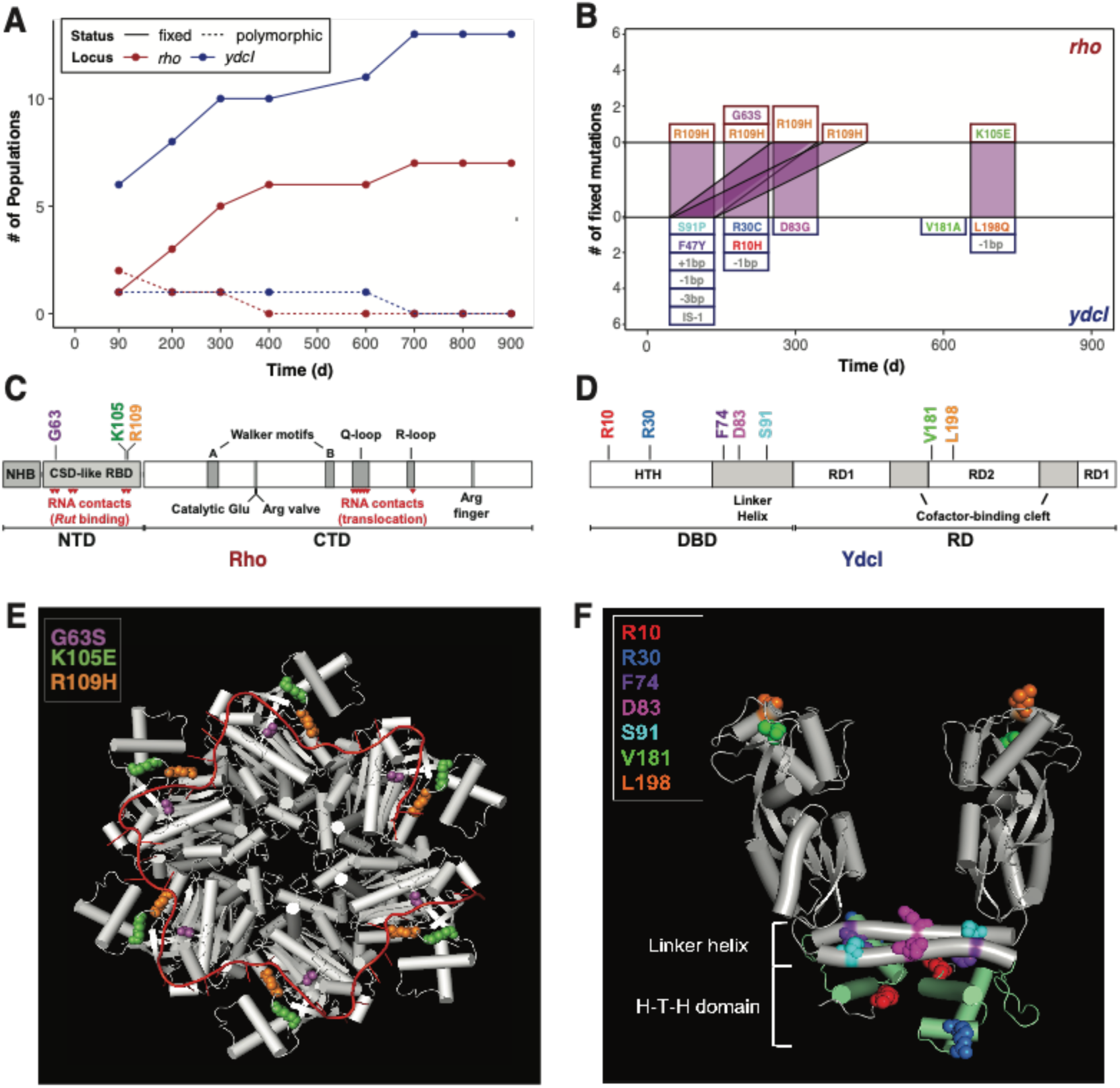
Repeated long-term starvation selects for mutations in *rho* and *ydcI*. **(A)** Number of populations containing polymorphic (dashed line) or fixed (solid line) mutations in the *rho* (red) or *ydcI* (blue) loci over 900 days of evolution to RLTS conditions. **(B)** Number of fixed mutations in *rho* (top half) and *ydcI* (bottom half) genes indicating timing at which each particular mutation arose over 900 days. Purple lines indicate alleles co-occurring within populations. The genic location of affected **(C)** *rho* (orange: R109H, purple: G63S, green: K105E) and **(D)** *ydcI* residues (red: R10H, blue: R30C, purple: F74Y, magenta: D83G, cyan: S91P, green: V181A, orange: L198Q) and **(E)** structural locations of affected *rho* and **(F)** *ydcI* residues identified in populations over 900 days of evolution to RLTS conditions. Only the A and B chains of the YdcI homo tetramer are visualized for simplicity. Structural locations colored in green indicate H-T-H domain residues. NTD: N-terminal domain, CTD: C-terminal domain, NHB: N-terminal helix bundle, CSD: cold shock domain, RBD: RNA binding domain, DBD: DNA binding domain, RD: regulatory domain, HTH: helix-turn-helix.

Since prior investigations implicate the evolution of arginine to histidine substitutions as a response to pH ^36,37,63,64^, and histidine residues can serve as pH sensors ^35,65,66^, we refocused our investigation to include genes shaped by positive selection during RTLS and associated with pH homeostasis. We found that all populations with mutant *rho* alleles also contain mutations in the recently annotated *ydcI* gene, which encodes a LysR-type transcriptional regulator reported to affect the expression of around 60 genes in *E. coli*, some of which are associated with the pH response ^67,68^ (**Fig 1A**; **Fig 1B; Fig S2B**). Mutations in *ydcI* are pervasive and evolved in the vast majority (13/16) of RTLS populations (**Table S1**). In each of these cases, the *ydcI* mutation arose simultaneously or before the *rho* mutation, suggesting that mutations in *ydcI* may potentiate the fixation of mutations in *rho*.

In contrast to the localized mutations identified in *rho*, the mutations in *ydcI* are diverse, ranging from putative loss of function mutations - caused by IS-element insertions and small indels - to nonsynonymous SNPs whose functional consequences on YdcI activity are currently unknown (**Fig 1D**). Nonsynonymous mutations in *ydcI* tend to cluster into three general genic regions. Using previous structural data for LysR-type proteins and the AlphaFold predicted structure for YdcI, we mapped the locations of these mutations to specific domains to gain insight into their potential functional impact (**Fig 1F**). The affected residues R10 and R30 are found in the predicted helix-turn-helix domain and could lead to an abolishment of DNA binding activity. Residues F74, D83, and S91 are all found in the α4 linker helix, and residues V181 and L198 are present in the junction between the two regulatory domains, regions that are critical to dimerization and effector binding, respectively, in well-studied LysR-type proteins ^69^. Together, this suggests that mutations resulting in loss or attenuation of YdcI function are under positive selection in RLTS conditions, and disruption of YdcI function may be necessary for the *rho* R109H allele to be beneficial.

### Rho^R109H^ mutants display a pH-sensitive in vitro phenotype

Previous investigations have identified arginine to histidine substitutions that alter protein functionality in alkaline conditions ^37,64^. Thus, we hypothesized that the activity of Rho^R109H^ may differ from that of WT Rho (Rho^+^) under alkaline conditions. To this end, we performed *in vitro* assays evaluating the capability of Rho to function as a helicase, terminate transcription, and bind RNA in neutral (pH 7) and alkaline (pH 9) conditions (**Fig 2; Fig S5**). At pH 7, we found no significant differences in the helicase activity (t-test; *P* = 0.829) or transcription termination patterns of Rho^R109H^ and only modest differences in RNA affinity (t-test, *P* = 0.033) compared to Rho^+^. By contrast, at pH 9, Rho^R109H^ exhibits significant reductions in its helicase activity (t-test; *P* = 5.54 x 10^-5^), a complete inability to terminate transcription, and a significant decline in RNA affinity (t-test, *P* = 0.003), while Rho^+^ remained functional. Moreover, Rho^+^ binding to RNA is largely unaffected by the change in pH conditions (t-test, *P* = 0.068), while there is a significant reduction in Rho^R109H^ RNA binding as pH increases (t-test, *P* = 0.004). These data collectively demonstrate that the activity of the Rho^R109H^ mutant is similar to the WT protein under neutral conditions, but is strongly reduced under alkaline conditions due, in part, to compromised RNA binding.

**Fig 2:**
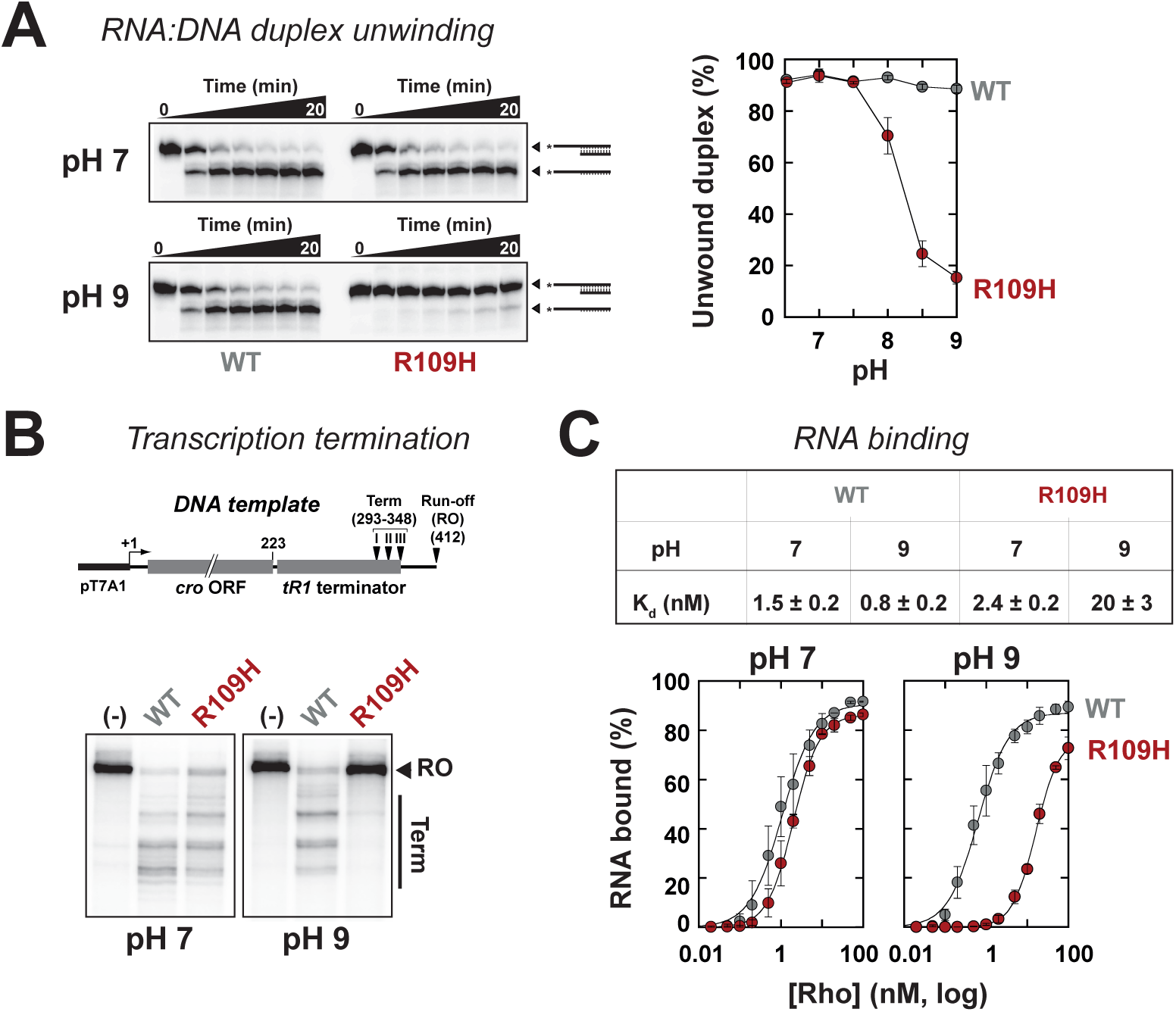
*In vitro* Rho Assay Reveals Diminished Activity of Rho^R109H^ at pH 9. The *in vitro* activities of Rho^+^ (WT) and Rho^R109H^ (R109H) were assessed at pH 7 and pH 9 in three ways: (**A**) efficiency of ATP-dependent unwinding of an RNA:DNA duplex containing a 5’-single-stranded *rut* overhang (graph: helicase activity after 20 min), (**B**) ability to terminate transcription at the λtR1 terminator, and (**C**) ability to bind to RNA. A schematic of the DNA template used in the transcription termination assay is shown above gel images in (**B**). Chart above plots in **(C)** contains the calculated average K_d_ for WT and R109H proteins. Additional information regarding binding parameters can be found in **Table 1**.

**Table 1:** Equilibrium binding parameters for Rho-RNA complexes.

| Parameters | WT |  | R109H |  |
| --- | --- | --- | --- | --- |
|  | pH 7 | pH 9 | pH 7 | pH 9 |
| n | 1.2 ± 0.2 | 1.3 ± 0.2 | 1.6 ± 0.2 | 1.7 ± 0.2 |
| K<sub>d</sub> (nM) | 1.5 ± 0.2 | 0.8 ± 0.2 | 2.4 ± 0.2 | 20 ± 3 |

<sup>a</sup> binding parameters were determined by non-linear least-squares fit of the slot-blot data to the Hill equation: $F_s = F_{max}[Rho]^n / ([Rho]^n + K_d^n)$ where $F_{max}$ is the maximal fraction of Rho-RNA complexes, $K_d$ is the dissociation constant, and $n$ is the Hill parameter reflecting binding cooperativity.

Of the transcripts where termination is Rho-dependent, a fraction of termination events require an additional transcription-termination factor, NusG ^43,45^, such as transcription termination that represses toxic prophage genes ^42^. These NusG-stimulated terminators have distinct *rut* features ^43,45^ and are less sensitive to two disruptive primary binding site mutations, Rho^F62S^ and Rho^Y80C^, than canonical terminators ^70^. To test the possibility that the R109H mutation may differentially impact the two classes of terminators, we conducted *in vitro* transcription experiments with two DNA templates encoding the NusG-stimulated terminators located within *aspS* and *yaiC* ^45^. With Rho^+^, termination at the *aspS* and *yaiC* sites is strongly stimulated by NusG at both pH 7 and pH 9 (**Fig S6**). Termination with the Rho^R109H^ mutant is also stimulated by NusG at pH 7, albeit to a lesser extent than Rho^+^. At pH 9, however, Rho^R109H^ can no longer trigger termination at the *aspS* and *yaiC* sites, even in the presence of NusG. These data indicate that pH regulates the activity of the Rho^R109H^ mutant at both NusG-stimulated and canonical Rho-dependent terminators (**Fig 2B**).

### Cells containing *rho* R109H/*ΔydcI* alleles have an altered intracellular pH response

The finding that Rho^R109H^ displays altered activity under alkaline conditions implies there must be changes in intracellular pH (pHi) to reveal the pH-dependent phenotype. Thus, we aimed to determine the pHi of cells containing these alleles using the dye BCECF-AM, commonly used to measure bacterial pHi ^71–74^. The intensity of the fluorescent signal emitted from BCECF-stained cells is dependent on pH ^73^, enabling the determination of the pHi of single cells when viewed under a fluorescent microscope. To ascertain if the introduction of the *rho* R109H and/or Δ*ydcI* alleles results in an alteration of pHi homeostasis, we determined the pHi of stained cells when incubated in a range of pH-adjusted buffers (pH 6.92 to 9.59) and compared their pHi response to that of WT cells.

Across the environmental pH conditions, WT cells exhibited pHi values with a fitted range from 6.7 to 9.05 (**Fig 3; Fig S8**), while *rho* R109H cells displayed a pHi range comparable to WT cells, 7.04 to 8.96 (**Fig 3; Fig S9**). Cells containing an *ydcI* deletion (Δ*ydcI*) exhibited a reduced pHi range, 7.48 to 8.92, with cells incubated in the lower pH range solutions yielding a higher pHi compared to WT cells (**Fig 3; Fig S10**). Importantly, *rho* R109H/Δ*ydcI* cells displayed a significant upshift of pHi compared to WT with a range of 7.53 to 9.67 (sign test; n=8; *P* = 0.008) (**Fig 3C; Fig S11**). Cells incubated in the more acidic pH solutions yielded an increased pHi most similar to that of Δ*ydcI* cells; however, the increased pHi of *rho* R109H/Δ*ydcI* cells extended through the entire range of pH-adjusted solutions. Thus, it appears that *rho* R109H/Δ*ydcI* cells have a higher baseline of pHi which would increase the likelihood of cells experiencing pHi conditions conducive to impaired Rho functioning.

**Fig 3:**
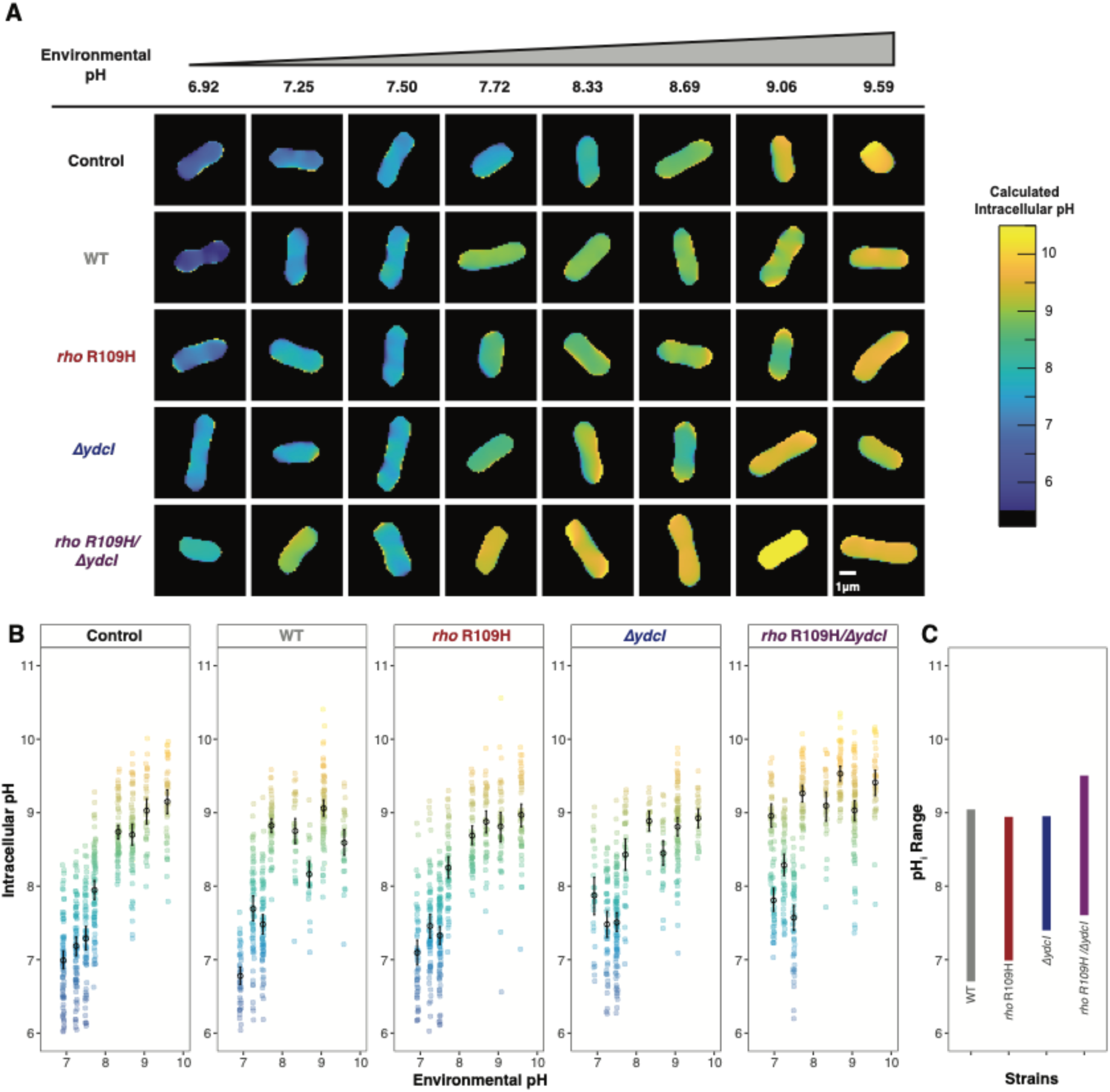
*rho* R109H/*ΔydcI* cells have increased intracellular pH. The intracellular pH (pHi) of WT, *rho* R109H, *ΔydcI*, and *rho* R109H/*ΔydcI* single cells were measured in buffers with increasing pH values. These measurements were derived from a ladder control (Control) containing WT cells treated with nigericin, yielding the pHi and environmental pH equal. (**A**). Representative microscopic images of single cells from tested strains in each pH buffered solution with colors of cells corresponding to their calculated pHi. Each image is cropped to 5.5 x 5.5 μm; a 1 μm scale bar is included in the bottom right image. Remaining microscopic images can be found in supplementary files. (**B**). pHi measurements derived from these images across the environmental pH conditions. Lightly shaded data points represent single cell replicates with the color denoting the pHi. Error bars display 95% confidence intervals. (**C**). Plot displaying the pHi range of each strain in tested conditions using data from panel (**B**) illustrates an upward shift in pHi range in the *rho* R109H/*ΔydcI* cells.

### Benefits of *rho* R109H and Δ*ydcI* alleles vary in RLTS environment

Considering the frequent co-occurrence of the *rho* R109H and Δ*ydcI* alleles in evolved populations and the necessity of both alleles for a significant alteration in pHi response, we hypothesized that the benefit of carrying the *rho* R109H allele could only be present in a Δ*ydcI* background. To address this, we “replayed” one cycle of the RLTS regime by growing cultures from different starting genotypes (WT, *rho* R109H, Δ*ydcI*) for 100 days and sequenced the *rho* and *ydcI* loci from the aged cultures (**Fig 4A; Fig S12**). We found that 8/12 populations beginning with a *rho* R109H genotype picked up putative Δ*ydcI* mutations, while *rho* R109H was present in 3/12 populations beginning with a Δ*ydcI* genotype. Importantly, we did not observe the emergence of the *rho* R109H allele in a population without detecting a putative Δ*ydcI* mutation. In fact, the only time we detected the *rho* R109H allele without a Δ*ydcI* was in 3/12 populations that began with the *rho R109H* genotype. Thus, *ydcI* loss-of-function mutations are recurrent in RLTS conditions and the appearance of the *rho* R109H mutation is unlikely to occur without a prior mutation in the *ydcI* gene.

**Fig 4:**
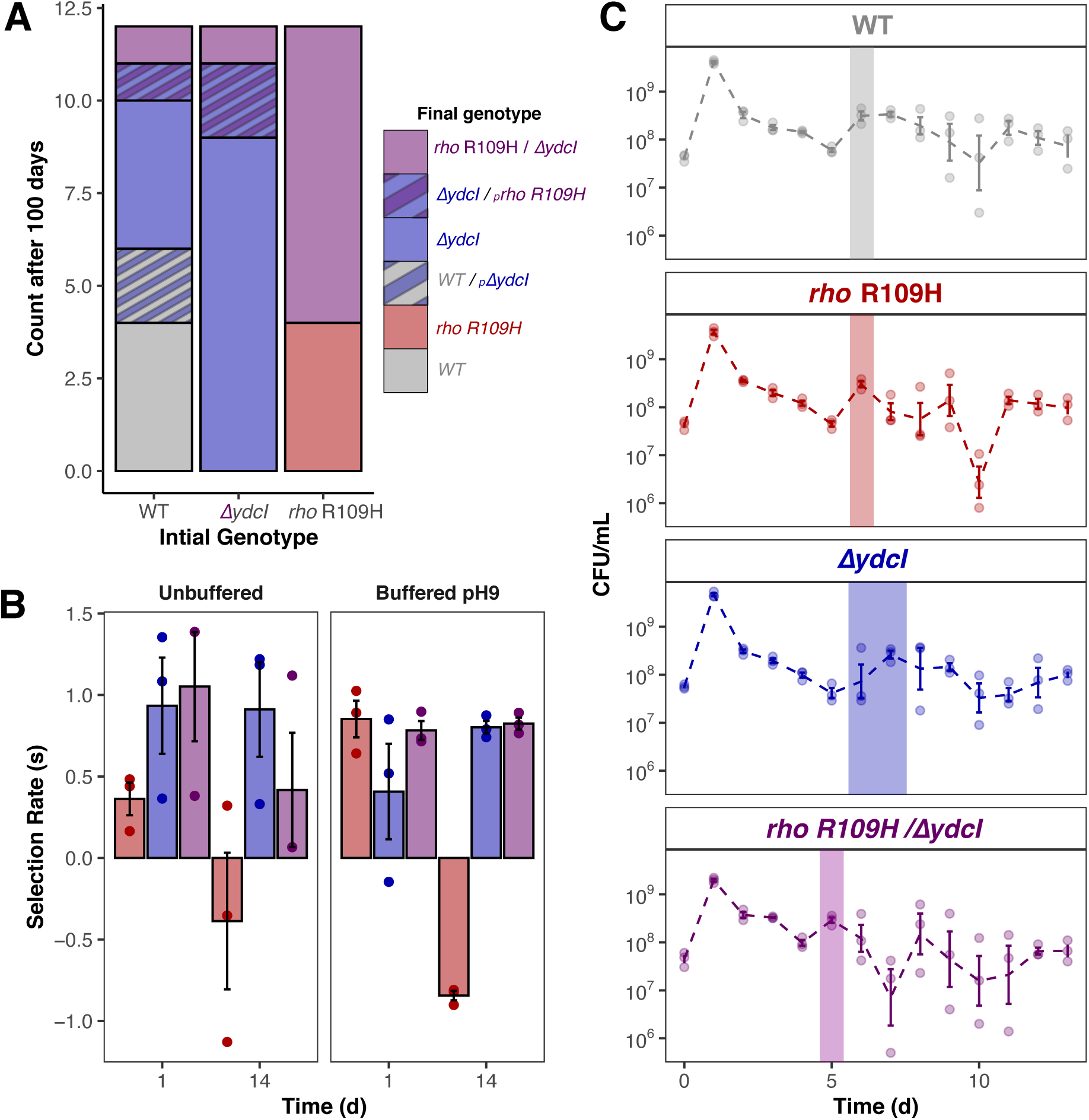
Benefits of *rho* R109H and *ΔydcI* alleles vary throughout the RLTS regime. **(A).** Frequency of final genotypes identified in populations after 12 replicate cultures of WT, *ΔydcI*, and *rho* R109H (initial genotypes) were grown in RLTS conditions for 100 days. A subscript of p denotes the allele is polymorphic as indicated by the Sanger sequence. **(B).** Pairwise competitions between WT ancestor and mutant strains when co-cultured for 1 or 14 days in unbuffered LB and LB buffered at pH 9. Colored bars indicate the median selection rate (s) for each mutant strain relative to the WT with single points representing each of three biological replicates. **(C).** Median CFU/mL of WT and mutant strains over 14 days of growth. Lightly shaded data points represent each of three biological replicates. Shaded vertical bars indicate the day of the first initial growth rebound post-death phase. Error bars display 95% confidence intervals.

Identifying the precise conditions under which each allele confers an advantage poses a challenge due to the continual evolution of the strains and the rapid accumulation of mutations as cultures age. While this is evident after “replaying” the evolution experiment, it may also account for the lack of significant differences in population cell density between strains over 100 days (**Fig S1B**). Consequently, we opted to conduct relevant phenotyping experiments over shorter periods to minimize any confounding effects potentially associated with the strains’ evolution. To mimic conditions experienced by populations during the RLTS regime, we performed pairwise competitions between each mutant and the WT ancestor in either unbuffered or buffered pH 9 LB medium for 1 and 14 days (**Fig 4B**). When competing the WT ancestor against the *rho* R109H strain, we observe an early benefit to carrying the mutant *rho* allele after the first day of co-culture, particularly in the pH 9 medium. However, after 14 days of co-culture the WT ancestor strongly outcompetes the *rho* R109H strain in both the unbuffered and pH 9 media. In contrast to *rho* R109H, Δ*ydcI* appears to exhibit considerably greater selection rates across the 14 days as the Δ*ydcI* strain outcompetes the WT ancestor at both time points and in both the unbuffered and buffered pH 9 media. Notably, when the *rho* R109H allele is present in a Δ*ydcI* background, we no longer observe negative selection rates after 14 days. Instead, the double *rho* R109H/Δ*ydcI* mutant strain outcompetes the WT strain under all conditions and maintains a high selection rate throughout the experiment. These results indicate that the *rho* R109H allele is advantageous when resources are plentiful, especially in alkaline conditions, and disrupting the *ydcI* gene can mediate any tradeoffs associated with the *rho* R109H allele in resource-limited long-term stationary phase conditions.

In addition to assessing growth and survival in co-culture, assessment of each strain in monoculture revealed no significant differences in the mutants’ survival over 14 days. The end of the stationary phase and subsequent onset of the death phase marks a considerable decline in population cell viability, which we consistently observe on day 3 of growth across all strains ^75^ (**Fig 4C; Fig S1B**). Next, the transition into long-term stationary phase is characterized by a resurgence in population density following the death phase, attributed to the substantial availability of dead cells providing resources for the surviving population ^75^. This resurgence in population cell density is highly-repeatable, occurring on the 6th day of growth for the WT ancestor, *rho* R109H, and Δ*ydcI* strains (**Fig 4C**). Notably, *rho* R109H/Δ*ydcI* populations exhibit a resurgence a day earlier on day 5 of growth. This early population resurgence following the death phase highlights a unique advantage of *rho* R109H/Δ*ydcI* as these cells can better utilize new resources as they become available, such as necromass and other metabolites released by dying cells.

### Sequence analysis identifies *rho* R109H alleles in natural isolates

Given the widespread presence of Rho across the bacterial domain and its essential role in many prokaryotes ^76^, we investigated the prevalence and distribution of a histidine residue at analogous positions within Rho (**Fig 5**). Our initial search of the RefSeq database did not return any bacterial sequences with histidine at this location. However, a deeper search using PHI-BLAST identified several sequences from species isolated from diverse environments that harbor a histidine residue at positions analogous to the *E. coli* Rho R109. Many of these isolates originated from alkaline environments or those which fluctuate between alkaline and neutral conditions. The soil bacterium *Inquilinus limosus* was isolated from a region that reports alkalization of the soil ^77,78^, while the *Poribacteria* genome was identified as part of a microbial mat in hypersaline, alkaline water ^79,80^. The pathogen *Bartonella baciliformis,* the causative agent of Carrion’s Disease, a neglected tropical disease, also encodes a histidine residue in Rho that may confer a pH-sensing phenotype as the pathogen alternates between neutral human blood and the alkaline midgut of sand flies ^25,81^. The remaining species are isolated from aquatic environments with reported pH gradients. Planctomyecetota was identified in sediment from the Auka marine hydrothermal vent (pH 6.5) in the Pacific Ocean (pH 8) ^82^, while Elusimicrobia was isolated from groundwater with a pH ranging from pH 6.5 - 8.5 ^83,84^. Together, the absence of a histidine residue in the analogous position 109 of Rho in various lab-adapted bacterial reference strains, as well as the identification of Rho H109 residues in isolates that experience alkaline environments, is consistent with our results showcasing that this allele provides beneficial effects in alkaline environments.

**Fig 5:**
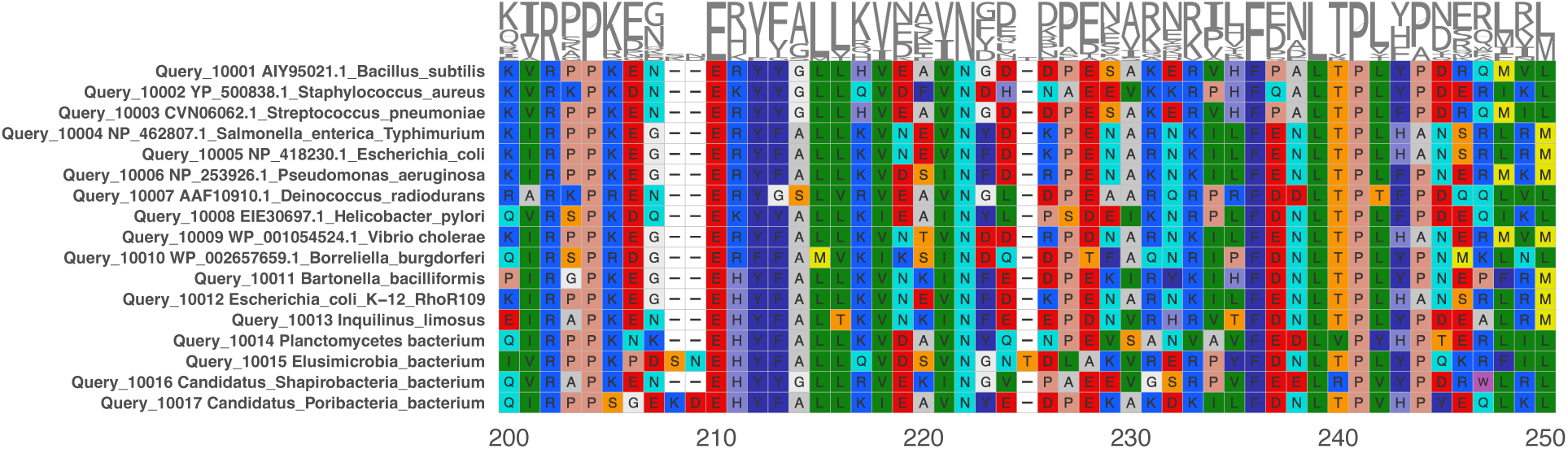
Amino acid alignment of Rho N-terminal RNA binding domain in laboratory strains and natural isolates. Alignments of Rho sequences from selected RefSeq genomes (top) and the mutated R109H *E. coli* clone with natural non-*E. coli* isolates (bottom) that encode a Histidine residue at the analogous position. Natural non-*E. coli* isolates were identified with PSI-BLAST.

## Discussion

Here, we report a novel adaptive pH-sensing mutation in the global transcriptional terminator Rho (*rho* R109H) that emerged independently in multiple *E. coli* populations subjected to 900 days of RLTS conditions. *In vitro* analyses revealed a pronounced pH-dependent loss of function, where Rho^R109H^ activity exhibited a sharp decline in the transition from neutral pH 7 to alkaline pH 9 conditions. This mutation was exclusively observed in strains that also contained a mutation in the transcriptional regulator gene *ydcI*, implying that in *E. coli* the *rho* R109H allele is beneficial in the RLTS environment when YdcI activity is attenuated or nonfunctional. Measurements of the pHi provide insight into the coexistence of these two alleles as we found that cells carrying both *rho* R109H and Δ*ydcI* alleles, exhibit a significantly more alkaline pHi, with respect to WT cells, across a spectrum of pH-adjusted solutions. These results imply that the intracellular environment of *rho* R109H/Δ*ydcI* cells is conducive to conditions where Rho^R109H^ termination would be impaired. Pairwise competitions between the mutants and WT ancestor pinpoint distinct advantages of each allele in the RTLS environment. Results of these competitions suggest that the *rho* R109H allele is beneficial during nutrient resource repletion as this mutation alone confers a fitness benefit only when resources are abundant, with an enhanced fitness when grown in LB broth adjusted to pH 9. However, this advantage is transient, as by 14 days of coculture, the *rho* R109H allele is no longer beneficial by itself. Instead, we observed that the deletion of *ydcI* masks the detrimental effects of the *rho* R109H allele during resource-depleted conditions, illustrating that the elimination of YdcI function generates an intracellular environment that better tolerates the plasticity of the *rho* R109H allele. Collectively, these data reveal a nuanced interplay between both alleles as they jointly provide unique advantages while navigating the complex stresses inherent to RLTS.

Prior research investigating the consequences of Rho impairment and the effects of various *rho* mutations on gene expression offers insight into the potential impacts of the *rho* R109H allele on resource utilization during pH fluctuations. In general, the impairment of Rho activity leads to an overall enrichment in transcripts due to transcriptional readthrough of Rho-dependent terminators ^85^. However, it appears that the identity and degree of differentially expressed genes varies based on the location and nature of the *rho* mutation, creating a challenge in pinpointing the exact benefit of the *rho* R109H allele. Although adaptive Rho mutations in residues that constitute the primary binding site are less common ^86,87^, these mutants do appear to be less disruptive to gene expression and have less enrichment in prophage gene expression, compared to evaluated secondary binding site mutants, which could partially explain why the *rho* R109H mutation is not detrimental ^42,70^. In some primary binding site mutants, a defect in RNA binding can be overcome at terminators with high NusG dependence ^70^. However, the Rho^R109H^ *in vitro* activity at two such terminators, *aspS* and *yaiC*, is also impaired in pH 9 conditions, signifying that the effects of this mutation vary from previously described primary binding site mutants. Adaptive mutations in *rho* can confer the ability to use novel substrates due to the upregulation of many metabolic pathways ^56,88^. Rho also controls the expression of genes related to the pH response ^42^, including tryptophanase, one of the most abundant proteins present in LB broth buffered to pH 9 ^89^. Adaptation to complex stresses elicits a distinct response that differs from the response to individual stresses ^90^. Thus, the pleiotropic nature of Rho may enable it to effectively navigate complex stress ^62^. Although it is currently unclear which genes are differentially expressed in *rho* R109H, it’s likely that changes in gene expression induced by this mutation in alkaline conditions impact both the pH response and metabolism. Moreover, since we do not observe fixation of the *rho* R109H mutation in cells without a deletion of *ydcI*, it is possible that the combination of these two alleles not only leads to changes associated with pHi, but also could relieve any detrimental effects on gene expression induced by loss of Rho activity, especially in the long-term stationary phase portion of the RLTS regime, and help to explain why YdcI function has to be eliminated in order for the *rho* R109H allele to be beneficial. This idea is also supported by the fact that YdcI has been implicated in pH homeostasis ^67,68^ and found to be mutated in *E. coli* strains adapted to highly alkaline conditions ^91^.

Adaptive nonsynonymous mutations in *rho* often do not confer immediate fitness advantages and typically necessitate additional mutations for the development of specific phenotypes ^53,56,92^. This occurrence is exemplified in the *rho* R109H mutant as its mutant state is likely not actualized unless the cellular environment is altered through the elimination of YdcI. Within *rho* R109H/Δ*ydcI* cells, the elevated pHi would likely result in a more uniform activity state for Rho since the majority of histidine residues would be deprotonated, effectively genetically assimilating the plastic *rho* R109H allele in its beneficial mutant state ^93^. Physiologically, the alkalinized pHi of *rho* R109H/Δ*ydcI* cells would help alleviate stress associated with maintaining pH homeostasis in an alkalized environment. In these conditions, pH homeostasis is dependent on the activity of electrogenic antiporters, such as the NhaA Na^+^/H^+^ pump, to power proton entry, placing a large energetic burden on maintaining a proton motive force ^24^. Upstream of *nhaA* lies *sokC*, which contains a Rho-dependent terminator ^43,94^. Thus, it is possible that when Rho activity is compromised, transcriptional readthrough from *sokC* leads to increased expression of *nhaA*. Moreover, by maintaining a pHi that is more alkaline than the environment, *rho* R109H/Δ*ydcI* cells would be less dependent on the activity of cation/proton antiporters to maintain homeostasis. This physiological benefit extends to the pH fluctuation period during resource replenishment when there is an abrupt downshift to a neutral environmental pH. In this condition, our data indicate that *rho* R109H/Δ*ydcI* cells would remain highly alkaline, leading to an even larger difference in protons across the membrane that would favor proton entry. Given that many metabolic substrates are often co-transported with H^+^ via symporters, the increased ΔpH of *rho* R109H/Δ*ydcI* cells would help to drive the import of resources ^95^. Thus, the alkalinization of *rho* R109H/Δ*ydcI* cells would not only help to reduce the energy required to maintain pH homeostasis, but would also enable the ability to better capitalize on resources through more efficient transport of the available resources.

In summary, these data report a novel pH-sensing mutation in an essential global regulator that confers significant physiological effects when paired with the inactivation of a mostly unknown transcription factor related to pH homeostasis. These mutations primarily evolve in the first 300 days of experimental evolution, illustrating how phenotypically plastic traits can facilitate rapid adaptation to complex, fluctuating environments. Despite the absence of this mutation from the current literature, we identify several natural isolates from fluctuating neutral/alkaline conditions carrying this allele, illustrating the power of experimental evolution for identifying functionally important mutations relevant to natural environments. Finally, the phenomenon described herein may signify genetic assimilation as a widespread adaptive mechanism for challenging complex environments. An emerging hallmark of cancer involves perturbations in pHi that manifest similarly to *rho* R109H/Δ*ydcI* cells, where the pHi is maintained higher than that of the extracellular environment ^96,97^. These alterations can be induced through arginine to histidine mutations that confer pH-sensing capabilities, particularly in global regulators, subsequently altering gene expression and reshaping the cellular environment ^36,37^. This suggests particular amino acid mutational signatures may yield effects applicable across all domains of life and that a greater understanding of how these patterns contribute to changes in cellular physiology is of paramount importance.

## Materials and Methods

### Strain construction and growth conditions

All bacterial strains originate from PFM2, a prototrophic derivative of *Escherichia coli* MG1655 K-12 that was provided by Dr. Patricia Foster (Indiana University) ^98^. The *rho* R109H substitution was cloned into the PFM2 *ΔaraBAD* background using the Church protocol ^99^. The *ΔydcI* and *rho* R109H/*ΔydcI* mutants were generated by moving the ydcI723(del)::kan cassette from the Keio collection strain JW5226 ^100^ to the PFM2 WT and *rho* R109H strains, respectively, using P1 vir transduction ^101^. Mutations were confirmed via PCR and Sanger sequencing. The *rho* R109H mutation was confirmed using a forward primer sequence of 5’-GGTGATGGCGTACTGGAGAT-3’ and a reverse primer sequence of 5’-GTGCCACAATCAGACCACG-3’. For the *ydcI* locus, mutations were confirmed using the forward primer 5’-CGGCACGATTAACTGAGGCCGT-3’ and reverse primer 5’-CGCCGTGATGCTGTTCGCCAA-3’.

### Identification of mutations in time-series whole-population metagenomic sequencing

Mutations in *rho* and *ydcI* were initially detected during the analysis of longitudinal metagenomic sequencing of 16 *E. coli* populations that were experimentally evolved in 100-day feast/famine cycle conditions ^18^. These populations originate from four different genetic backgrounds: PFM2; PFM2, *ΔaraBAD;* PFM2*, ΔmutL; or* PFM2*, ΔaraBAD, ΔmutL.* The deletion of the *ΔaraBAD l*ocus serves as a neutral marker that enables the detection of cross-contamination between lines during experimental evolution by plating on TA agar (10 g/L tryptone, 1g/L yeast extract, 5 g/L NaCl, 16 g/L agar, 10 g/L L-arabinose, 0.005% tetrazolium chloride) ^102^, while the deletion of *ΔmutL* disrupts methyl-directed mismatch repair and allows for the examination of the evolutionary process in a mutator phenotype. The relevant genotypes of populations adapted to 100-day feast/famine cycles can be found in **Table S1**.

### Structural Analysis

The quaternary structure of YdcI was predicted with SWISS-MODEL from an alpha-fold predicted structure for YdcI (P77171) on UniprotKB. As YdcI is annotated as a LysR-family transcriptional regulator which are typically homo-tetramers ^103,104^, we focused our search on the homology options in the SWISS-MODEL repository that were homo-tetramers and based on OxyR -another member of the LysR family (templates: 4×6g.1.A; and 6g1d.1.A). Residues that acquired nonsynonymous mutations throughout experimental evolution were highlighted in PyMol. For the structure of Rho, the most recent and most detailed cryoEM reconstruction of an *E. coli* Rho-RNA complex was used (#8E6W), while Pymol was used to highlight features of interest.

### In vitro analysis of Rho activity

Plasmid pET28bR109H for overexpression of the R109H mutant was obtained by site-directed mutagenesis of plasmid pET28bRho (kindly provided by Pr. James Berger, Johns Hopkins University) using the NEBuilder HiFi kit (New England Biolabs). The R109H mutant was overexpressed in BL21(DE3)pLysS cells carrying the pET28bR109H plasmid and purified as described for WT Rho ^105^.

Duplex unwinding experiments were performed as described ^105^ with minor modifications. Briefly, 5nM RNA:DNA duplexes bearing a ^32^P label and a *rut* site (substrate C ^106^) were incubated for 5 min at 37°C with 20 nM Rho hexamers in helicase buffer (150 mM potassium acetate and 20 mM of either MES, HEPES, or EPPS at the indicated pH). Then, 1 mM MgCl_2_, 1 mM ATP, and 400 nM Trap oligonucleotide (complementary to the DNA strand of the duplexes) were added to the reaction mixture before incubation at 30°C. Reaction aliquots were withdrawn at various times, quenched with two volumes of quench buffer (20 mM EDTA, 0.5% SDS, 150 mM sodium acetate, 4% Ficoll-400), and analyzed by 9% polyacrylamide gel electrophoresis (PAGE) and phosphorimaging (Typhoon FLA-9500 instrument and ImageQuantTL v8.1 software, Cytivia).

Transcription termination experiments were performed as described ^107^ with minor modifications. Briefly, DNA template encoding the λtR1 terminator (0.1 pmol), *E. coli* RNA polymerase (0.4 pmol; New England Biolabs), Rho (1.4 pmol), and Superase-In (0.5U/µL; Ambion) were mixed in 18 µL of transcription buffer (150 mM potassium acetate, 5 mM MgCl_2_, 1mM DTT, 0.05 mg/mL BSA and either 20 mM HEPES, pH 7.0 or EPPS, pH 9.0) and incubated for 10 min at 37°C. Then, 2 µL of initiation mix (2 mM ATP, GTP, and CTP, 0.2 mM UTP, 2.5 µCi/µL of ^32^P-aUTP, 250 µg/mL rifampicin) were added to the mixtures before further incubation for 20 min at 37°C. Reactions were quenched with 70 mM EDTA, 0.1mg/mL tRNA, and 0.3 M sodium acetate, phenol extracted, concentrated by ethanol precipitation, and then analyzed by denaturing 8% PAGE and phosphorimaging.

Rho-RNA dissociation constants (K_d_) were determined using a filter-binding assay, as described ^108^. Briefly, ∼10 fmoles of ^32^P-labeled RNA substrate (RNA strand from substrate C) were mixed with various amounts of Rho in 100 µL of binding buffer (0.1 mM EDTA, 1 mM DTT, 150 mM potassium acetate, 20 µg/mL BSA, and either 20 mM HEPES, pH 7.0; or 20 mM EPPS, pH 9.0). After incubation for 10 min at 37°C, the samples were filtered through stacked [top] nitrocellulose (Amersham Protran) and [bottom] cationic nylon (Pall Biodyne B) membranes using a Bio-dot SF apparatus (Biorad). The fractions of free and Rho-bound RNA (retained on the nylon and nitrocellulose membranes, respectively) as a function of Rho concentration (expressed in hexamers) were then determined by phosphorimaging of the membranes.

### Quantification of intracellular pH

Intracellular pH was determined using the pH-sensitive fluorescent dye BCECF-AM. Cultures of each experimental strain, including an additional WT culture used for calibration of radiometric fluorescence calibration (control), were grown overnight in LB broth. The following day, a 1 mL aliquot from each culture was washed two times via centrifugation and resuspended in 500 µL of unbuffered Ringer’s buffer (1/4X) (Oxoid). We stained cells for 30 minutes at 37C with 2 µM BCECF-AM (Invitrogen). In addition to BCECF-AM staining, we added 5 µL of Nigericin (Invitrogen) to the tube containing the WT ladder standard to allow intracellular pH to equilibrate with the external environment. Following staining, we performed two additional washes via centrifugation using 500 µL of unbuffered 1/4X Ringers’ solution (Oxoid). 100 µL aliquots of each experimental strain and the ladder sample were transferred into five separate tubes. We centrifuged each tube for 1 minute at 13,000 x g and resuspended the pellets in one of seven 1/4X buffered Ringers’ solutions (pH 6.92, 7.25, 7.72, 8.33, 8.69, 9.06, 9.59). We prepared buffered Ringers’ solutions using 50 mM of 1,3-Bis[tris(hydroxymethyl)amino] propane (Acros Organics) and adjusting the pH to the appropriate level using 1 M HCl. Once resuspended into pH-adjusted solutions, we imaged cells after approximately 30 minutes.

For each condition, 7 µL of cells were sandwiched between a 22 mm square #1.5 coverslip and a standard glass slide, providing sufficient fluid volume for cells to remain in solution but in approximately the same focal plane of the microscope. Cells were imaged on a Nikon Ti inverted microscope (Nikon Instruments, Melville, NY) equipped with an Andor Zyla sCMOS camera (Oxford Instruments, Abingdon, Oxfordshire), Spectra/Aura light engine (Lumencor, Beaverton, OR), and an Apo TIRF 60X Oil DIC N2 NA 1.49 objective (Nikon Instruments, Melville, NY). The BCECF fluorescence intensity was imaged in two channels (1: Ex 470 nm, Em 525/50 nm and 2: Ex 440 nm, Em 525/50 nm). We used a semiautomated MATLAB script (MathWorks, Natwick, MA) to segment cells and quantify the fluorescence intensity. In brief, images underwent a spatial bandpass filter (spatial frequencies between 220 nm and 1100 nm) to remove salt and pepper noise and background intensity. Objects were then segmented as individual regions if above a certain intensity threshold. Because cells were plated at a sufficiently low density to ensure single objects, a relative threshold for each image was set as the intensity such that approximately the top-1000 pixel values were above the threshold (quantile of 0.9998). To ensure that the entire region of each cell was included, and to deal with the fact that these two imaging channels have a small spatial shift from one channel to the other, these regions were then dilated by disks of radius 2 µm. Background regions were defined by dilating these regions by a further 2 µm, and removing the objects from these masks, yielding regions that are near, but not overlapping with, individual objects. The intensity for each object in each channel was determined as the background-subtracted sum of all the pixels in each region. Lists of relative intensities of these objects were then imported into Rstudio. Samples treated with nigericin were used to determine the coefficients A and B in a fit of pH_obs_ = A*log(intensityRatio)+B.

### Replaying of RLTS Regime

To determine the prevalence of the *rho* R109H and *ΔydcI* alleles in the RLTS condition and to parse out any patterns in their co-occurrence, we replayed one cycle of the RLTS regime by incubating WT, *rho* R109H, and *ΔydcI* cultures for 100 days after which time we examined the *rho* and *ydcI* loci for acquired mutations. We inoculated 12 cultures of WT, *rho* R109H, and *ΔydcI* strains in 10 mL of LB broth each originating from a separate colony from an LB agar plate freshly prepared from frozen glycerol stocks. We incubated each culture at 37°C and shaken upright at 180 rpm for a total of 100 days. We left cultures undisturbed during this time except for a brief period every 10 days when the culture tubes were gently removed from the incubator and added sterile deionized water to achieve a volume of 10 mL to counteract evaporation. After 100 days, we removed the cultures and centrifuged them 2500 x g for 10 minutes. We froze the resulting pellet of bacterial biomass in liquid nitrogen and it was stored at −80°C until DNA extraction was performed. We extracted DNA from the pellets using a DNeasy UltraClean Microbial Kit (Qiagen) and used the resulting DNA samples as template for PCR. We performed PCR using the same primers previously mentioned that were used for confirmation of the initial mutation. The PCR products were purified using the Monarch PCR & DNA Cleanup Kit (NEB) and sent for Sanger sequencing (GENEWIZ).

### Pairwise Competition Assays

To determine the fitness outcomes of the mutant strains relative to the WT ancestral strain, we performed pairwise competition assays ^102,109^. Here, culture tubes containing a 50:50 starting proportion of two competing strains are grown for a given period, at which time they are sampled to determine the resulting proportions of each strain following growth. Each competing strain contains either an intact *araBAD* operon or *ΔaraBAD* genotype and can be differentiated from each other by plating on TA agar (10 g/L tryptone, 1 g/L yeast extract, 5 g/L NaCl, 16 g/L agar, 10 g/L L-arabinose, 0.005% tetrazolium chloride) where the WT *araBAD* strain produces light pink colonies and the *ΔaraBAD* strain produces bright red colonies. Overnight cultures were prepared for each strain using colonies picked from freshly streaked LB agar plates obtained from frozen stocks. For each competition, we inoculated 10 mL of LB broth with 50 µL of overnight culture from both competitors. We set up individual competitions for each sampling time point, as vortexing and resampling of the same competition tube has been shown to alter fitness outcomes ^23^. Before incubation, we sampled each competition to determine precise starting proportions by collecting a 100 µL aliquot, serially diluting in PBS, and plating on TA agar. All competition tubes were incubated at 37°C until the sampling time point at which time a 100 µL sample was removed, serially diluted in PBS, and plated on TA agar. All TA agar plates were incubated at 37°C for 24 hours before colonies were enumerated.

### Survival Assays

We assessed the survival of our strains by monitoring their population density over the course of 14 days. We inoculated 3 replicate overnight cultures of WT, *rho* R109H, *ΔydcI,* and *rho* R109H/*ΔydcI* strains in 10 mL of LB broth each originating from a separate colony from an LB agar plate freshly prepared from frozen glycerol stocks and incubated them for 18 hours at 37°C. The survival assay was initiated by aliquoting 100 µL of each overnight culture into 14 individual culture tubes filled with 10 mL of fresh LB broth. We incubated each culture at 37°C, shaken upright at 180 rpm until the appropriate sampling time point. We set up individual cultures for each sampling time point, as vortexing and resampling of the same culture tube has been shown to alter fitness outcomes ^23^. Every 24 hours, the culture tube corresponding to the sampling time point was removed from the incubator and gently vortexed. We then removed 100 µL from the culture and performed serial dilutions in PBS. We plated 100 µL aliquots of the dilutions on LB agar plates and enumerated the colonies from countable plates after 24 hours of incubation at 37°C.

### Identification of Other Bacterial *rho* R109H Alleles

Rho homologs containing a histidine at the analogous *Escherichia coli* R109 residue were identified using blastp with the Pattern Hit Initiated (PHI-BLAST) algorithm ^110^. The following PHI pattern EH[YF][YFG][AGS][LM][LVT] targets R109H substitutions to the conserved ERFYALL motif from the *Escherichia coli* Rho sequence (NP_418230.1) while allowing for any variation observed after alignment to Rho sequences with COBALT ^111^ across nine additional species of bacteria spanning the bacterial tree of life: Bacillus subtilis (AIY95021.1); *Borreliella burgdorferi* (WP_002657659.1); *Deinococcus radiodurans* (AAF10910.1); *Helicobacter pylori* (EIE30697.1); *Pseudomonas aeruginosa* (NP_253926.1); *Salmonella enterica* typhimurium (NP_462807.1); *Staphylococcus aureus* (YP_500838.1); *Streptococcus pneumoniae* (CVN06062.1); and *Vibrio cholerae* (WP_001054524.1) (**Fig 5**).

## Acknowledgements

We would like to thank the Advanced Computing Center for Research and Education (ACCRE) and Vanderbilt Cell Imaging Shared Resource (supported by NIH grants CA68485, DK20593, DK58404, DK59637 and EY08126) specifically the Nikon Multi-Excitation TIRF Microscope acquired under S10-OD018075-01A1.

This work was supported by Army Research Office grant W911NF-21-1-0161 (M.G.B.), National Institutes of Health grant R35GM150625 (M.G.B.), and French Agence Nationale de la Recherche grant ANR-19-CE44-0009-01 (M.B.) Additional funds were provided by Vanderbilt University (startup funds: M.G.B.), Vanderbilt University Medical Center (startup funds: B.P.B), Evolutionary Studies Initiative at Vanderbilt (M.G.B and B.P.B), and the VUMC Discovery Scholars in Health and Medicine Program (B.P.B).

## Author Contributions

S.B. Worthan: Methodology, Formal Analysis, Investigation, Writing-original draft, Writing-review and editing, Visualization; R.D.P. McCarthy: Methodology, Formal Analysis, Investigation, Writing-original draft, Visualization; M. Delaleau: Methodology, Formal Analysis, Investigation, Visualization; R. Stikeleather: Methodology, Resources; B. P. Bratton: Methodology, Software, Formal Analysis, Investigation, Writing-review and editing, Visualization; M. Boudvillain: Methodology, Formal Analysis, Investigation, Writing-review and editing, Visualization, Funding acquisition; M.G. Behringer: Methodology, Software, Conceptualization, Formal Analysis, Investigation, Writing-original draft, Writing-review and editing, Visualization, Funding Acquisition

## Notes

### Competing Interest Statement

The authors have declared no competing interest.

## References

1. Aertsen, A. & Michiels, C. W. Stress and how bacteria cope with death and survival. Crit Rev Microbiol 30, 263–273 (2004).

2. Lambert, G. & Kussell, E. Memory and Fitness Optimization of Bacteria under Fluctuating Environments. PLOS Genetics 10, e1004556 (2014).

3. Nguyen, J., Lara-Gutiérrez, J. & Stocker, R. Environmental fluctuations and their effects on microbial communities, populations and individuals. FEMS Microbiology Reviews 45, (2021).

4. Morita, R. Y. Bioavailability of energy and its relationship to growth and starvation survival in nature. Can. J. Microbiol. 34, 436–441 (1988).

5. Reese, A. T. et al. Microbial nitrogen limitation in the mammalian large intestine. Nat Microbiol 3, 1441–1450 (2018).

6. Hobbie, J. & Hobbie, E. Microbes in nature are limited by carbon and energy: the starving-survival lifestyle in soil and consequences for estimating microbial rates. Front. microbiol 4, (2013).

7. Kolter, R., Siegele, D. A. & Tormo, A. The Stationary Phase of the Bacterial Life Cycle. Annu. Rev. Microbiol. 47, 855–874 (1993).

8. Zambrano, M. M., Siegele, D. A., Almirón, M., Tormo, A. & Kolter, R. Microbial Competition: Escherichia coli Mutants That Take Over Stationary Phase Cultures. Science 259, 1757–1760 (1993).

9. Zambrano, M. M. & Kolter, R. GASPing for life in stationary phase. Cell 86, 181–184 (1996).

10. Finkel, S. E. Long-term survival during stationary phase: evolution and the GASP phenotype. Nat Rev Microbiol 4, 113–120 (2006).

11. Katz, S. et al. Dynamics of Adaptation During Three Years of Evolution Under Long-Term Stationary Phase. Molecular Biology and Evolution 38, 2778–2790 (2021).

12. Ratib, N. R., Seidl, F., Ehrenreich, I. M. & Finkel, S. E. Evolution in Long-Term Stationary-Phase Batch Culture: Emergence of Divergent Escherichia coli Lineages over 1,200 Days. mBio 12, e03337–20 (2021).

13. Shoemaker, W. R. et al. Microbial population dynamics and evolutionary outcomes under extreme energy limitation. PNAS 118, e2101691118 (2021).

14. Aouizerat, T. et al. Eukaryotic Adaptation to Years-Long Starvation Resembles that of Bacteria. iScience 19, 545–558 (2019).

15. Behringer, M. G., et al. Escherichia coli cultures maintain stable subpopulation structure during long-term evolution. PNAS 115, E4642–E4650 (2018).

16. Behringer, M. G. et al. Complex Ecotype Dynamics Evolve in Response to Fluctuating Resources. mBio 13, e03467–21 (2022).

17. Kram, K. E., et al. Adaptation of Escherichia coli to Long-Term Serial Passage in Complex Medium: Evidence of Parallel Evolution. mSystems 2, 10.1128/msystems.00192-16 (2017).

18. Behringer, M. G. et al. Trade-offs, trade-ups, and high mutational parallelism underlie microbial adaptation to extreme feast/famine. 2023.10.04.560893 Preprint at 10.1101/2023.10.04.560893 (2023).

19. Ratzke, C. & Gore, J. Modifying and reacting to the environmental pH can drive bacterial interactions. PLoS Biol 16, e2004248 (2018).

20. Farrell, M. J. & Finkel, S. E. The Growth Advantage in Stationary-Phase Phenotype Conferred by rpoS Mutations Is Dependent on the pH and Nutrient Environment. J Bacteriol 185, 7044–7052 (2003).

21. Sezonov, G., Joseleau-Petit, D. & D’Ari, R. Escherichia coli Physiology in Luria-Bertani Broth. J Bacteriol 189, 8746–8749 (2007).

22. Sánchez-Clemente, R., Guijo, M. I., Nogales, J. & Blasco, R. Carbon Source Influence on Extracellular pH Changes along Bacterial Cell-Growth. Genes 11, 1292 (2020).

23. Worthan, S. B., McCarthy, R. D. P. & Behringer, M. G. Case Studies in the Assessment of Microbial Fitness: Seemingly Subtle Changes Can Have Major Effects on Phenotypic Outcomes. J Mol Evol 91, 311–324 (2023).

24. Krulwich, T. A., Sachs, G. & Padan, E. Molecular aspects of bacterial pH sensing and homeostasis. Nat Rev Microbiol 9, 330–343 (2011).

25. Santos, V. C., Araujo, R. N., Machado, L. A. D., Pereira, M. H. & Gontijo, N. F. The physiology of the midgut of Lutzomyia longipalpis (Lutz and Neiva 1912): pH in different physiological conditions and mechanisms involved in its control. Journal of Experimental Biology 211, 2792–2798 (2008).

26. Lande, R. Evolution of phenotypic plasticity and environmental tolerance of a labile quantitative character in a fluctuating environment. Journal of Evolutionary Biology 27, 866–875 (2014).

27. O’Brien, P. A., Morrow, K. M., Willis, B. L. & Bourne, D. G. Implications of Ocean Acidification for Marine Microorganisms from the Free-Living to the Host-Associated. Front. Mar. Sci. 3, (2016).

28. Yi, X. & Dean, A. M. Phenotypic plasticity as an adaptation to a functional trade-off. eLife 5, e19307 (2016).

29. Hoskisson, P. A. & Hutchings, M. I. MtrAB–LpqB: a conserved three-component system in actinobacteria? Trends in Microbiology 14, 444–449 (2006).

30. Rebek, J. On the structure of histidine and its role in enzyme active sites. Struct Chem 1, 129–131 (1990).

31. Rötzschke, O., Lau, J. M., Hofstätter, M., Falk, K. & Strominger, J. L. A pH-sensitive histidine residue as control element for ligand release from HLA-DR molecules. PNAS 99, 16946–16950 (2002).

32. Williamson, D. M., Elferich, J. & Shinde, U. Mechanism of Fine-tuning pH Sensors in Proprotein Convertases: Identification of a ph-sensing histidine pair in the propeptide of proprotein convertase 1/3. J Biol Chem 290, 23214–23225 (2015).

33. Vercoulen, Y. et al. A Histidine pH sensor regulates activation of the Ras-specific guanine nucleotide exchange factor RasGRP1. eLife 6, e29002 (2017).

34. Gerchman, Y. et al. Histidine-226 is part of the pH sensor of NhaA, a Na+/H+ antiporter in Escherichia coli. Proc. Natl. Acad. Sci. U.S.A. 90, 1212–1216 (1993).

35. Webb, B. A., et al. A Histidine Cluster in the Cytoplasmic Domain of the Na-H Exchanger NHE1 Confers pH-sensitive Phospholipid Binding and Regulates Transporter Activity. JBC 291, 24096–24104 (2016).

36. Szpiech, Z. A. et al. Prominent features of the amino acid mutation landscape in cancer. PLOS ONE 12, e0183273 (2017).

37. White, K. A. et al. Cancer-associated arginine-to-histidine mutations confer a gain in pH sensing to mutant proteins. Sci Signal 10, eaam9931 (2017).

38. Elena, S. F. & Lenski, R. E. Evolution experiments with microorganisms: the dynamics and genetic bases of adaptation. Nat Rev Genet 4, 457–469 (2003).

39. McDonald, M. J. Microbial Experimental Evolution – a proving ground for evolutionary theory and a tool for discovery. EMBO Rep 20, e46992 (2019).

40. Saxer, G. et al. Mutations in global regulators lead to metabolic selection during adaptation to complex environments. PLoS Genet 10, e1004872 (2014).

41. Fraebel, D. T., Gowda, K., Mani, M. & Kuehn, S. Evolution of Generalists by Phenotypic Plasticity. iScience 23, 101678 (2020).

42. Cardinale, C. J. et al. Termination Factor Rho and Its Cofactors NusA and NusG Silence Foreign DNA in E. coli. Science 320, 935–938 (2008).

43. Peters, J. M. et al. Rho and NusG suppress pervasive antisense transcription in Escherichia coli. Genes Dev 26, 2621–2633 (2012).

44. Sedlyarova, N. et al. sRNA-Mediated Control of Transcription Termination in E. coli. Cell 167, 111–121.e13 (2016).

45. Delaleau, M., Eveno, E., Simon, I., Schwartz, A. & Boudvillain, M. A scalable framework for the discovery of functional helicase substrates and helicase-driven regulatory switches. PNAS 119, e2209608119 (2022).

46. Washburn, R. S. & Gottesman, M. E. Transcription termination maintains chromosome integrity. PNAS 108, 792–797 (2011).

47. Kriner, M. A., Sevostyanova, A. & Groisman, E. A. Learning from the Leaders: Gene Regulation by the Transcription Termination Factor Rho. TIBS 41, 690–699 (2016).

48. Bossi, L., Figueroa-Bossi, N., Bouloc, P. & Boudvillain, M. Regulatory interplay between small RNAs and transcription termination factor Rho. BBA-Gene Regulatory Mechanisms 1863, 194546 (2020).

49. Steinmetz, E. J. & Platt, T. Evidence supporting a tethered tracking model for helicase activity of Escherichia coli Rho factor. PNAS 91, 1401–1405 (1994).

50. Molodtsov, V., Wang, C., Firlar, E., Kaelber, J. T. & Ebright, R. H. Structural basis of Rho-dependent transcription termination. Nature 614, 367–374 (2023).

51. Kishimoto, T. et al. Transition from Positive to Neutral in Mutation Fixation along with Continuing Rising Fitness in Thermal Adaptive Evolution. PLOS Genetics 6, e1001164 (2010).

52. Tenaillon, O. et al. The Molecular Diversity of Adaptive Convergence. Science 335, 457–461 (2012).

53. Rodríguez-Verdugo, A., Carrillo-Cisneros, D., González-González, A., Gaut, B. S. & Bennett, A. F. Different tradeoffs result from alternate genetic adaptations to a common environment. PNAS 111, 12121–12126 (2014).

54. Deatherage, D. E., Kepner, J. L., Bennett, A. F., Lenski, R. E. & Barrick, J. E. Specificity of genome evolution in experimental populations of Escherichia coli evolved at different temperatures. PNAS 114, E1904–E1912 (2017).

55. Boyd, S. M. et al. Experimental Evolution of Copper Resistance in Escherichia coli Produces Evolutionary Trade-Offs in the Antibiotics Chloramphenicol, Bacitracin, and Sulfonamide. Antibiotics 11, 711 (2022).

56. Freddolino, P. L., Goodarzi, H. & Tavazoie, S. Fitness Landscape Transformation through a Single Amino Acid Change in the Rho Terminator. PLOS Genetics 8, e1002744 (2012).

57. Haft, R. J. F. et al. Correcting direct effects of ethanol on translation and transcription machinery confers ethanol tolerance in bacteria. PNAS 111, E2576–2585 (2014).

58. Conrad, T. M. et al. Whole-genome resequencing of Escherichia coli K-12 MG1655 undergoing short-term laboratory evolution in lactate minimal media reveals flexible selection of adaptive mutations. Genome Biology 10, R118 (2009).

59. Herron, M. D. & Doebeli, M. Parallel Evolutionary Dynamics of Adaptive Diversification in Escherichia coli. PLOS Biology 11, e1001490 (2013).

60. Wolfe, A. J. The Acetate Switch. Microbiol Mol Biol Rev 69, 12–50 (2005).

61. Cesar, S. et al. Bacterial Evolution in High-Osmolarity Environments. mBio 11, e01191–20 (2020).

62. Mitra, P., Ghosh, G., Hafeezunnisa, M. & Sen, R. Rho Protein: Roles and Mechanisms. Annu Rev Microbiol 71, 687–709 (2017).

63. Damaghi, M., Wojtkowiak, J. W. & Gillies, R. J. pH sensing and regulation in cancer. Frontiers in Physiology 4, (2013).

64. DiGiammarino, E. L. et al. A novel mechanism of tumorigenesis involving pH-dependent destabilization of a mutant p53 tetramer. Nat Struct Mol Biol 9, 12–16 (2002).

65. Frantz, C. et al. Cofilin is a pH sensor for actin free barbed end formation: role of phosphoinositide binding. JCB 183, 865–879 (2008).

66. Choi, C.-H., Webb, B. A., Chimenti, M. S., Jacobson, M. P. & Barber, D. L. pH sensing by FAK-His58 regulates focal adhesion remodeling. JCB 202, 849–859 (2013).

67. Gao, Y. et al. Systematic discovery of uncharacterized transcription factors in Escherichia coli K-12 MG1655. Nucleic Acids Res 46, 10682–10696 (2018).

68. Romiyo, V. & Wilson, J. W. Phenotypes, transcriptome, and novel biofilm formation associated with the ydcI gene. Antonie van Leeuwenhoek 113, 1109–1122 (2020).

69. Maddocks, S. E. & Oyston, P. C. F. Structure and function of the LysR-type transcriptional regulator (LTTR) family proteins. Microbiology 154, 3609–3623 (2008).

70. Shashni, R., Qayyum, M. Z., Vishalini, V., Dey, D. & Sen, R. Redundancy of primary RNA-binding functions of the bacterial transcription terminator Rho. Nucleic Acids Res 42, 9677–9690 (2014).

71. Tsujimoto, K., Semadeni, M., Huflejt, M. & Packer, L. Intracellular pH of halobacteria can be determined by the fluorescent dye 2’, 7’-bis(carboxyethyl)-5(6)-carboxyfluorescein. BBRC 155, 123–129 (1988).

72. Molenaar, D., Bolhuis, H., Abee, T., Poolman, B. & Konings, W. N. The efflux of a fluorescent probe is catalyzed by an ATP-driven extrusion system in Lactococcus lactis. J Bacteriol 174, 3118–3124 (1992).

73. Riondet, C., Cachon, R., Waché, Y., Alcaraz, G. & Diviès, C. Measurement of the intracellular pH in Escherichia coli with the internally conjugated fluorescent probe 5- (and 6-)carboxyfluorescein succinimidyl ester. Biotechnol. Tech. 11, 735–738 (1997).

74. Heo, J., Thomas, K. J., Seong, G. H. & Crooks, R. M. A Microfluidic Bioreactor Based on Hydrogel-Entrapped E. coli: Cell Viability, Lysis, and Intracellular Enzyme Reactions. Anal. Chem. 75, 22–26 (2003).

75. Zinser, E. R. & Kolter, R. Mutations Enhancing Amino Acid Catabolism Confer a Growth Advantage in Stationary Phase. J. Bacteriol. 181, 5800–5807 (1999).

76. Banerjee, S., Chalissery, J., Bandey, I. & Sen, R. Rho-dependent Transcription Termination: More Questions than Answers. J Microbiol 44, 11–22 (2006).

77. Festa, S., Coppotelli, B. M. & Morelli, I. S. Bacterial diversity and functional interactions between bacterial strains from a phenanthrene-degrading consortium obtained from a chronically contaminated-soil. International Biodeterioration & Biodegradation 85, 42–51 (2013).

78. Ileana, P. et al. Soil Quality Problems Associated with Horticulture in the Southern Urban and Peri-Urban Area of Buenos Aires, Argentina. in Urban Horticulture - Necessity of the Future (IntechOpen, 2020). doi:10.5772/intechopen.90351.

79. Ruvindy, R., White III, R. A., Neilan, B. A. & Burns, B. P. Unravelling core microbial metabolisms in the hypersaline microbial mats of Shark Bay using high-throughput metagenomics. ISME J 10, 183– 196 (2016).

80. White, R. A., Wong, H. L., Ruvindy, R., Neilan, B. A. & Burns, B. P. Viral Communities of Shark Bay Modern Stromatolites. Front. microbiol 9, (2018).

81. Dostálová, A. & Volf, P. Leishmania development in sand flies: parasite-vector interactions overview. Parasites & Vectors 5, 276 (2012).

82. Speth, D. R. et al. Microbial communities of Auka hydrothermal sediments shed light on vent biogeography and the evolutionary history of thermophily. The ISME Journal 16, 1750–1764 (2022).

83. Hama, K., Kunimaru, T., Metcalfe, R. & Martin, A. J. The hydrogeochemistry of argillaceous rock formations at the Horonobe URL site, Japan. *Physics and Chemistry of the Earth*, Parts A/B/C 32, 170–180 (2007).

84. Hernsdorf, A. W. et al. Potential for microbial H2 and metal transformations associated with novel bacteria and archaea in deep terrestrial subsurface sediments. The ISME Journal 11, 1915–1929 (2017).

85. Peters, J. M. et al. Rho directs widespread termination of intragenic and stable RNA transcription. Proceedings of the National Academy of Sciences 106, 15406–15411 (2009).

86. Mohamed, E. T. et al. Generation of a platform strain for ionic liquid tolerance using adaptive laboratory evolution. Microbial Cell Factories 16, 204 (2017).

87. Du, B. et al. Adaptive laboratory evolution of Escherichia coli under acid stress. Microbiology (Reading) 166, 141–148 (2020).

88. Hafeezunnisa, Md. & Sen, R. The Rho-Dependent Transcription Termination Is Involved in Broad-Spectrum Antibiotic Susceptibility in Escherichia coli. Frontiers in Microbiology 11, (2020).

89. Stancik, L. M. et al. pH-dependent expression of periplasmic proteins and amino acid catabolism in Escherichia coli. J Bacteriol 184, 4246–4258 (2002).

90. Gunasekera, T. S., Csonka, L. N. & Paliy, O. Genome-Wide Transcriptional Responses of Escherichia coli K-12 to Continuous Osmotic and Heat Stresses. J Bacteriol 190, 3712–3720 (2008).

91. Hamdallah, I. et al. Experimental Evolution of Escherichia coli K-12 at High pH and with RpoS Induction. Appl Environ Microbiol 84, e00520–18 (2018).

92. Hug, S. M. & Gaut, B. S. The phenotypic signature of adaptation to thermal stress in Escherichia coli. BMC Evol Biol 15, 177 (2015).

93. Waddington, C. H. Genetic Assimilation of an Acquired Character. Evolution 7, 118–126 (1953).

94. Hafeezunnisa, M., Chhakchhuak, P. I. R., Krishnakumar, J. & Sen, R. Rho-dependent transcription termination regulates the toxin–antitoxin modules of cryptic prophages to silence their expression in Escherichia coli. FEBS Letters 595, 2057–2067 (2021).

95. Elferink, M. G. L., Hellingwerf, K. J. & Konings, W. N. The role of the proton motive force and electron flow in solute transport in Escherichia coli. European Journal of Biochemistry 153, 161–165 (1985).

96. Sharma, M. et al. pH Gradient Reversal: An Emerging Hallmark of Cancers. Recent Pat Anticancer Drug Discov 10, 244–258 (2015).

97. Koltai, T. et al. Chapter 5 - How pH deregulation favors the hallmarks of cancer. in pH Deregulation as the Eleventh Hallmark of Cancer (eds. Koltai, T. et al.) 101–121 (Academic Press, 2023). doi:10.1016/B978-0-443-15461-4.00008-4.

98. Lee, H., Popodi, E., Tang, H. & Foster, P. L. Rate and molecular spectrum of spontaneous mutations in the bacterium Escherichia coli as determined by whole-genome sequencing. PNAS 109, E2774–2783 (2012).

99. Mosberg, J. A., Gregg, C. J., Lajoie, M. J., Wang, H. H. & Church, G. M. Improving Lambda Red Genome Engineering in Escherichia coli via Rational Removal of Endogenous Nucleases. PLOS ONE 7, e44638 (2012).

100. Baba, T. et al. Construction of Escherichia coli K-12 in-frame, single-gene knockout mutants: the Keio collection. Mol Syst Biol 2, 2006.0008 (2006).

101. Saragliadis, A., Trunk, T. & Leo, J. C. Producing Gene Deletions in Escherichia coli by P1 Transduction with Excisable Antibiotic Resistance Cassettes. JoVE 58267 (2018) doi:10.3791/58267.

102. Levin, B. R., Stewart, F. M. & Chao, L. Resource-Limited Growth, Competition, and Predation: A Model and Experimental Studies with Bacteria and Bacteriophage. The American Naturalist 111, 3– 24 (1977).

103. Schell, M. A. Molecular biology of the LysR family of transcriptional regulators. Annu Rev Microbiol 47, 597–626 (1993).

104. Giannopoulou, E.-A. et al. Crystal structure of the full-length LysR-type transcription regulator CbnR in complex with promoter DNA. The FEBS Journal 288, 4560–4575 (2021).

105. Boudvillain, M., Walmacq, C., Schwartz, A. & Jacquinot, F. Simple Enzymatic Assays for the In Vitro Motor Activity of Transcription Termination Factor Rho from Escherichia coli. in Helicases: Methods and Protocols (ed. Abdelhaleem, M. M.) 137–154 (Humana Press, Totowa, NJ, 2010). doi:10.1007/978-1-60327-355-8_10.

106. Soares, E., Schwartz, A., Nollmann, M., Margeat, E. & Boudvillain, M. The RNA-mediated, asymmetric ring regulatory mechanism of the transcription termination Rho helicase decrypted by time-resolved nucleotide analog interference probing (trNAIP). Nucleic Acids Res 42, 9270–9284 (2014).

107. Nadiras, C., Eveno, E., Schwartz, A., Figueroa-Bossi, N. & Boudvillain, M. A multivariate prediction model for Rho-dependent termination of transcription. Nucleic Acids Res 46, 8245–8260 (2018).

108. D’Heygère, F., Schwartz, A., Coste, F., Castaing, B. & Boudvillain, M. ATP-dependent motor activity of the transcription termination factor Rho from Mycobacterium tuberculosis. Nucleic Acids Res 43, 6099–6111 (2015).

109. Lenski, R. E., Rose, M. R., Simpson, S. C. & Tadler, S. C. Long-Term Experimental Evolution in Escherichia coli. I. Adaptation and Divergence During 2,000 Generations. The American Naturalist 138, 1315–1341 (1991).

110. Zhang, Z. et al. Protein sequence similarity searches using patterns as seeds. Nucleic Acids Research 26, 3986–3990 (1998).

111. Papadopoulos, J. S. & Agarwala, R. COBALT: constraint-based alignment tool for multiple protein sequences. Bioinformatics 23, 1073–1079 (2007).

