## Supplemental Material for "Evolution of pH-sensitive transcription termination during adaptation to repeated long-term starvation"

**Supplemental Methods**

***Long-Term Survival Assays and Culture pH***

We assessed long-term survival of the WT, *rho* R109H, *ΔydcI*, and *rho* R109H/*ΔydcI* strains through periodic quantification of population cellular density via viable plate counts. We inoculated 3 replicate overnight cultures of WT, *rho* R109H, *ΔydcI,* and *rho* R109H/*ΔydcI* strains in 10 mL of LB broth each originating from a separate colony from an LB agar plate freshly prepared from frozen glycerol stocks and incubated them for 18 hours at 37°C. The long-term survival assay was initiated by aliquoting 100 µL of each overnight culture into 16 individual culture tubes filled with 10 mL of fresh LB broth with each tube representing a sampling time point. We incubated the cultures at 37°C, shaken upright at 180 rpm until the appropriate sampling time point. We set up individual cultures for each sampling time point, as vortexing and resampling of the same culture tube has been shown to alter fitness outcomes. We left cultures undisturbed during incubation except for a brief period every 10 days when the culture tubes were gently removed from the incubator and sterile diH_2_O was added to achieve a volume of 10 mL in efforts to counteract evaporation. We performed viable cell plate counts at time 0 by sampling the tubes immediately following inoculation. Thereafter, the population density was sampled after 1, 4, and 7 days, followed by every 7 days until 100 days. At the appropriate sampling time point, the corresponding culture tubes were removed from the incubator and gently vortexed. We then removed 100 µL from the cultures and performed serial dilutions in PBS. We plated a 100 µL aliquots of the dilutions on LB agar plates and enumerated the colonies from countable plates after 24 hours of incubation at 37°C. We also removed 1 mL aliquots from the culture tubes during sampling and removed the cells via centrifugation at 2500 x g for 5 minutes. We then measured the pH of the resulting supernatant using the Thermo Scientific Orion Star A214 pH meter.

***Growth Analyses***

We assessed growth of the *rho* R109H strain in conditions that previously produced other experimentally evolved *rho* mutations, including temperature stress, exposure to ethanol, and osmotic stress. All growth analyses were performed in 96-well plates and we used a Synergy H1 Plate Reader (Biotek) to analyze growth. A volume of 150 µL of LB broth was inoculated with 1.5 µL of overnight culture in each well so that every strain had three biological replicates. The 96-well plate underwent an 18 hour program at 37 °C with continuous orbital shaking at 807 cpm. Initial and final absorbance (600 nm) readings were taken, as well as intermittent absorbance readings every 15 min. Temperature stress was evaluated by growing the strains at three different temperatures: 32 °C, 37 °C, and 42 °C. Exposure to ethanol was measured by supplementing ethanol into LB broth using three different concentrations: 4%, 6%, and 7%. Finally, osmotic stress was evaluated in two ways: by supplementing cultures with NaCl  (0.1 M NaCl, 0.2 M NaCl, 0.5 M NaCl, and 1.0 M NaCl) and by supplementing cultures with sucrose (0.1 M sucrose, 0.2 M sucrose, 0.5 M sucrose, and 1.0 M sucrose). Growth curve parameters and statistics were calculated with the R package Growthcurver (v.0.3.1) which fits growth data to a logistic model.

**Supplemental Figures**

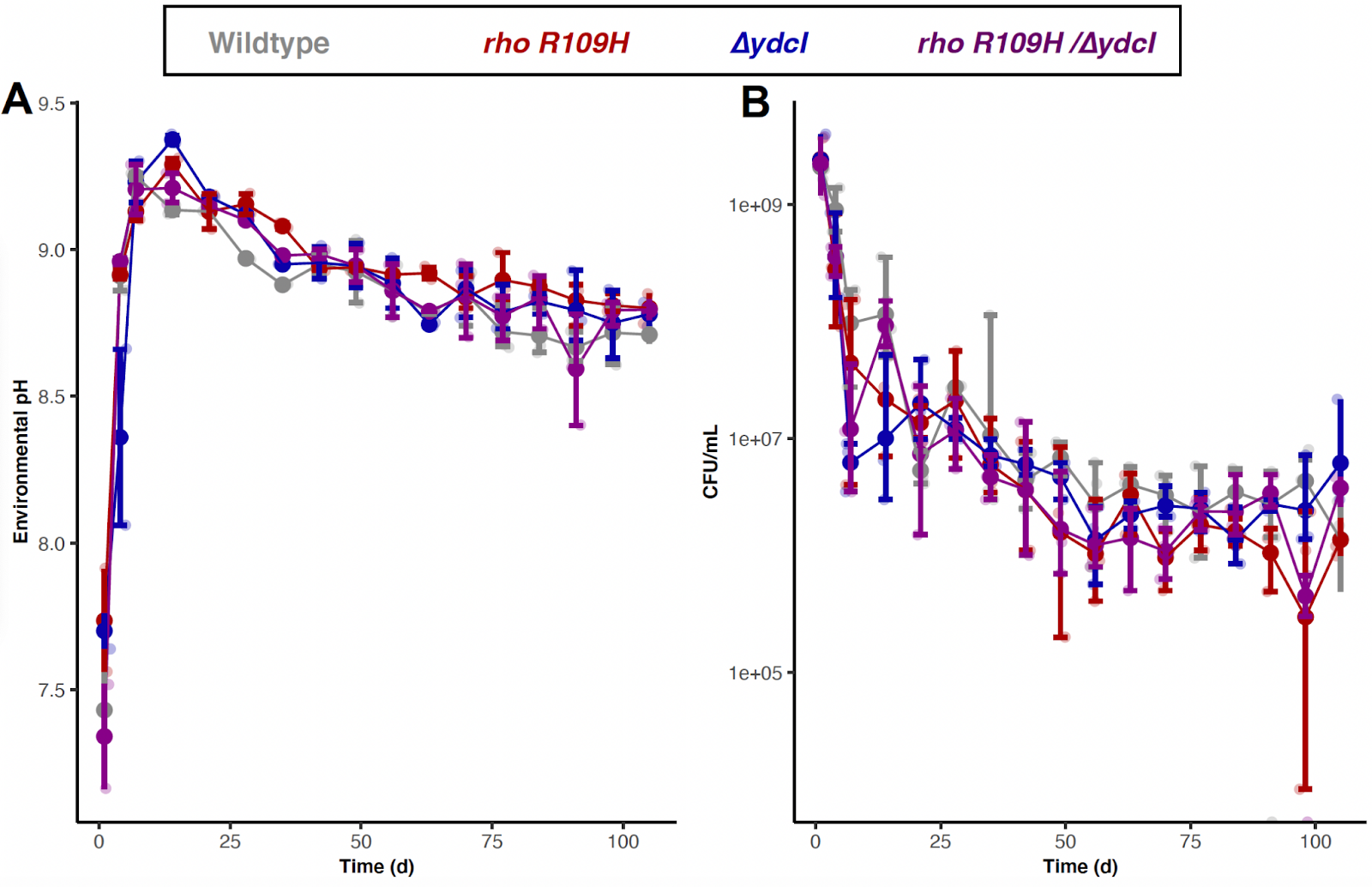

**Fig S1: Culture pH and Survival over 100 days. (A).** The medium pH (environmental pH) of cultures containing WT, *rho* R109H, *ΔydcI*, and *rho* R109H/*ΔydcI* were determined over the course of 100 days. **(B).** Survival of cultures, as measured by viable plate counts (CFU/mL), containing WT, *rho* R109H, *ΔydcI*, and *rho* R109H/*ΔydcI* were determined over the course of 100 days. Each data point represents the median of 3 biological replicate cultures. Error bars denote 95% confidence intervals.

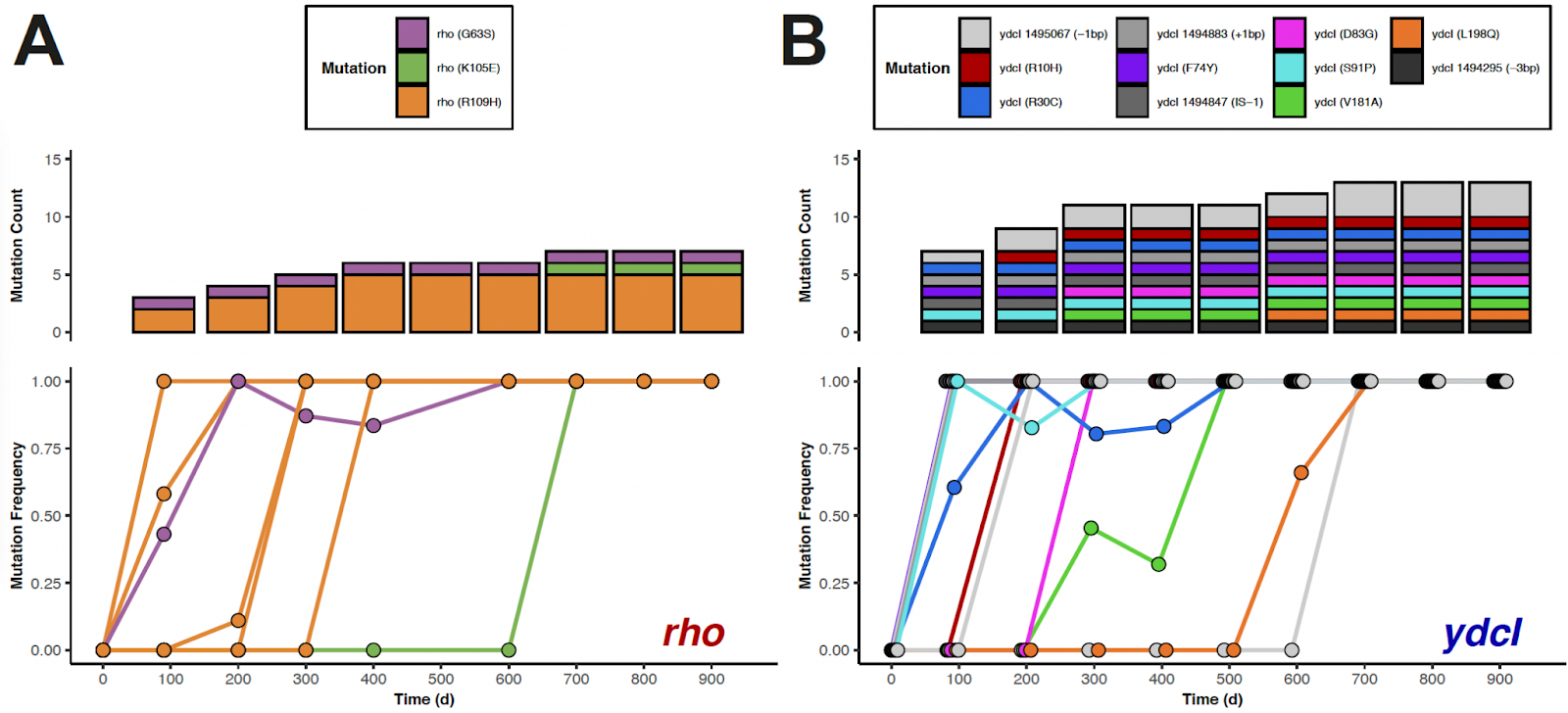

**Fig S2: Mutation dynamics in *rho* and *ydcI* over 900 days of experimental evolution.** The number (mutation count) and frequency of mutations in (**A**). *rho* and (**B**). *ydcI* genes in experimentally evolved populations over 900 days of RLTS. Colors in plots correspond to different alleles defined in the legends.

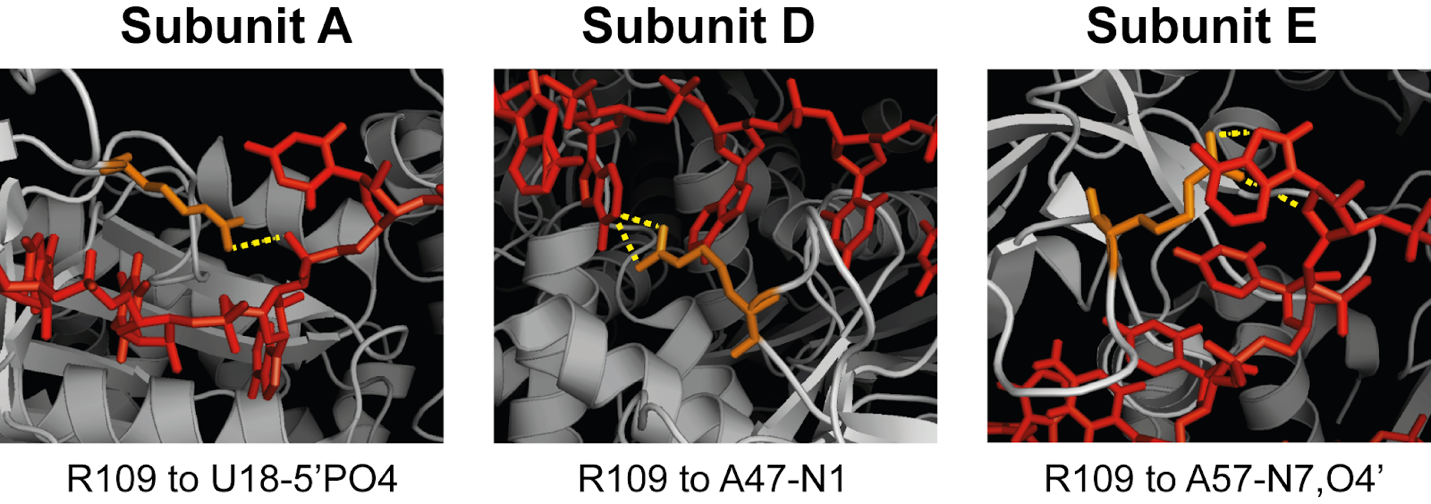

**Fig S3: Rho primary RNA binding site highlighting position of residue R109**. Direct R109 (orange) to RNA (red) contacts, indicated in yellow, in Rho subunits observed in PDB# 8E6W clearly illustrates R109 as part of the RNA primary binding site.

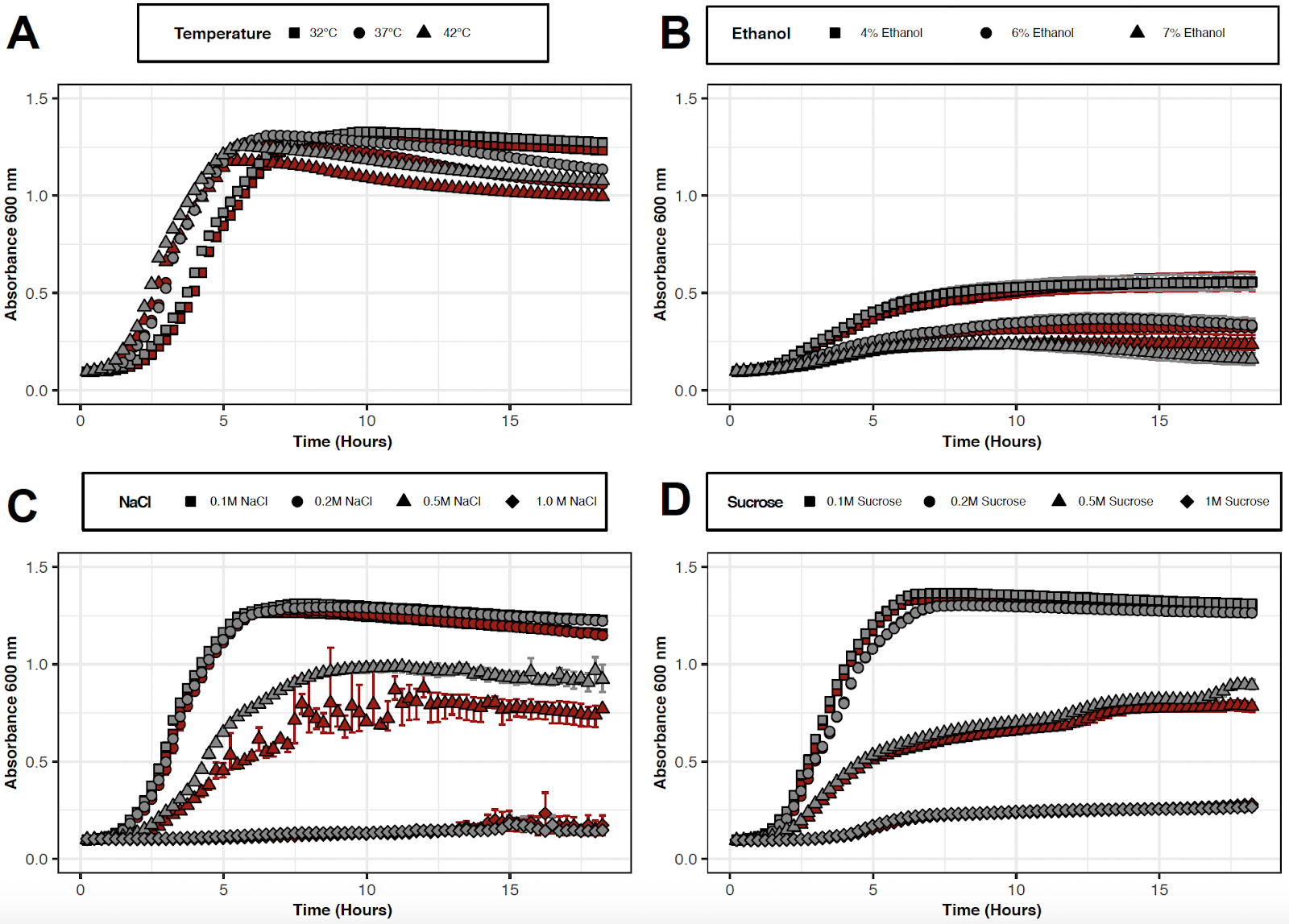

**Fig S4: Growth Analyses of *rho* R109H in conditions that previously produced other experimentally evolved *rho* mutations.** WT (gray) and *rho* R109H (red) cultures were grown for 18 hours in LB broth at varying (**A**). temperatures or supplemented with several concentrations of (**B**). ethanol, (**C**). NaCl, and (**D**). sucrose. Each data point represents the median of three biological replicates. Error bars represent 95% confidence intervals.

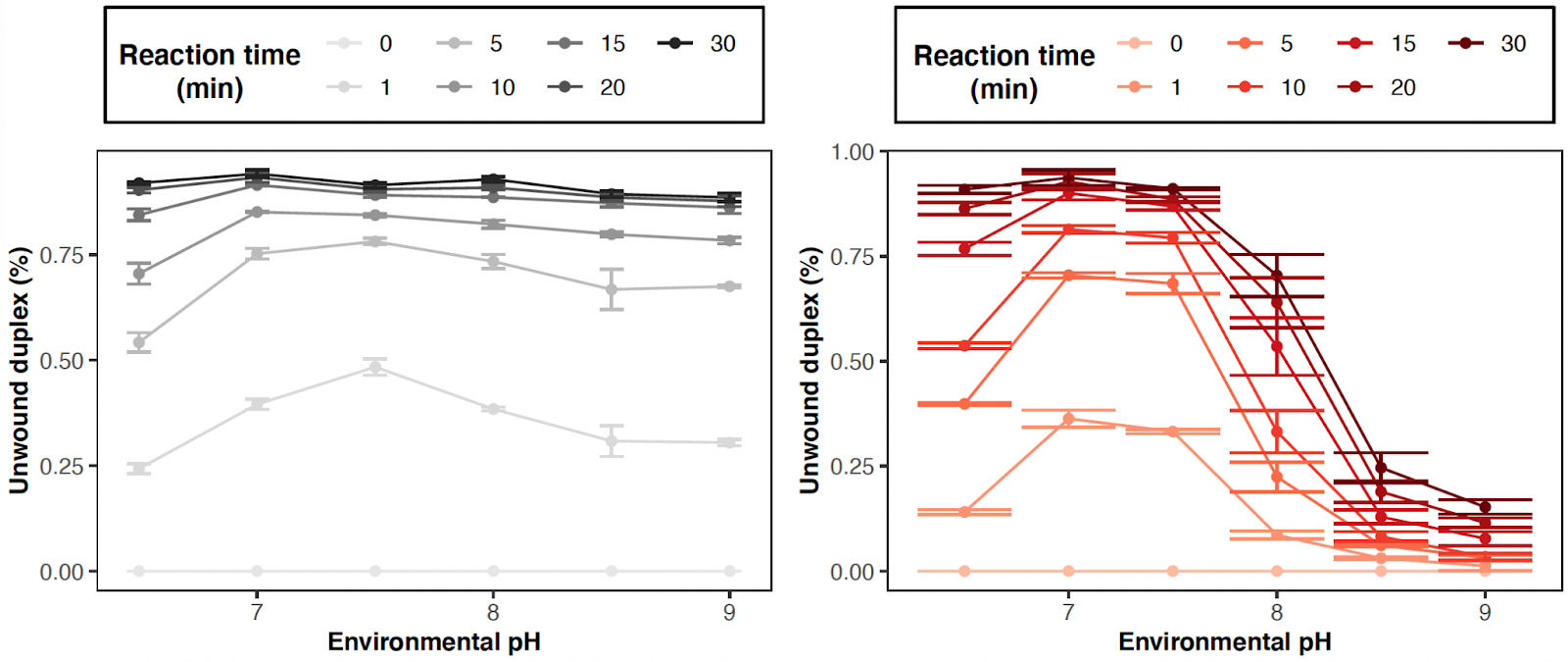

**Fig S5: Helicase activity of Rho to unwind a RNA:DNA duplex.** Percent of unwound duplex over the reaction time (min) across increasing environmental pH conditions for Rho^+^ (gray scale) and Rho^R109H^ (red scale) proteins. Each data point represents the median of 2 biological replicates with error bars representing 95% confidence intervals.

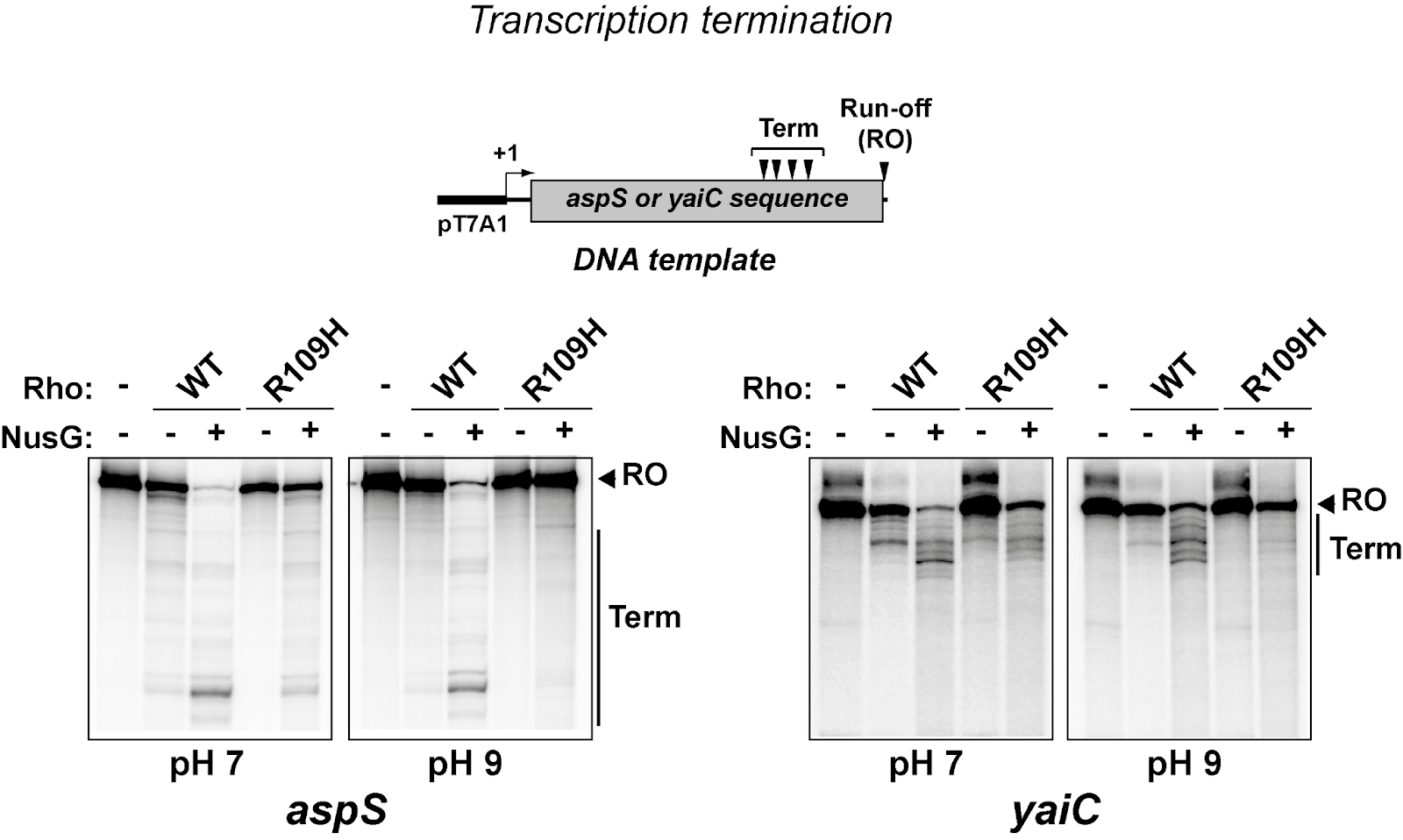

**Fig S6: Termination with the R109H mutant is also dependent on pH at NusG-stimulated terminators.** Transcription termination *in vitro* activity of Rho^+^ (WT) and Rho^R109H^ (R109H) proteins at NusG-stimulated terminators in *aspS* and *yaiC* genes in pH 7 and pH 9 conditions. Schematic above gel images denotes the design of DNA template used for the termination activity assay.

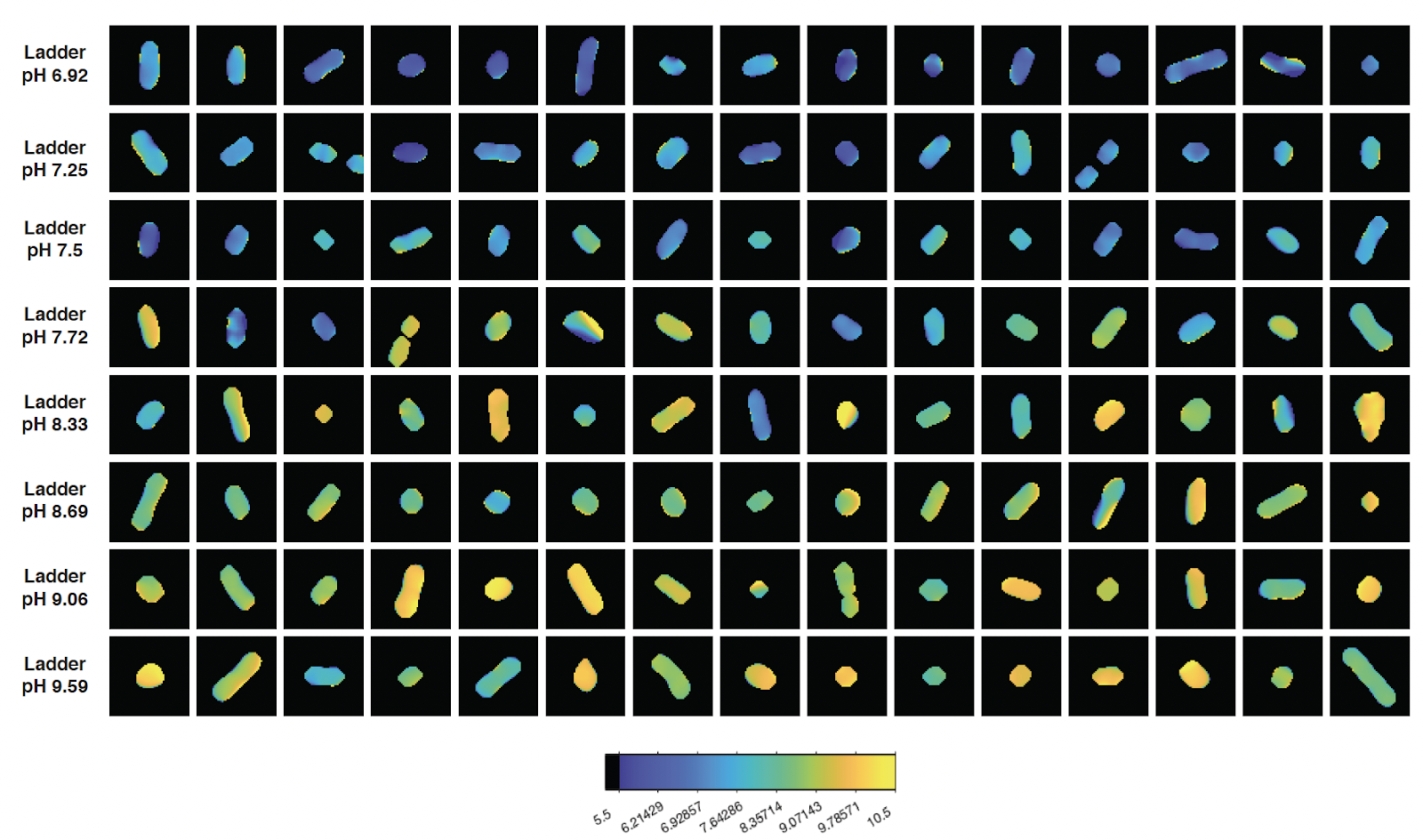

**Fig S7: Control Ladder for measurement of intracellular pH.** Microscopic images of all replicate WT cells treated with nigericin, yielding the intracellular and environmental pH equal, used to produce calibration curve. Color of cells correspond to the cellular pH as denoted on the scale below microscopic images. Each image is cropped to 5.5 x 5.5 μm.

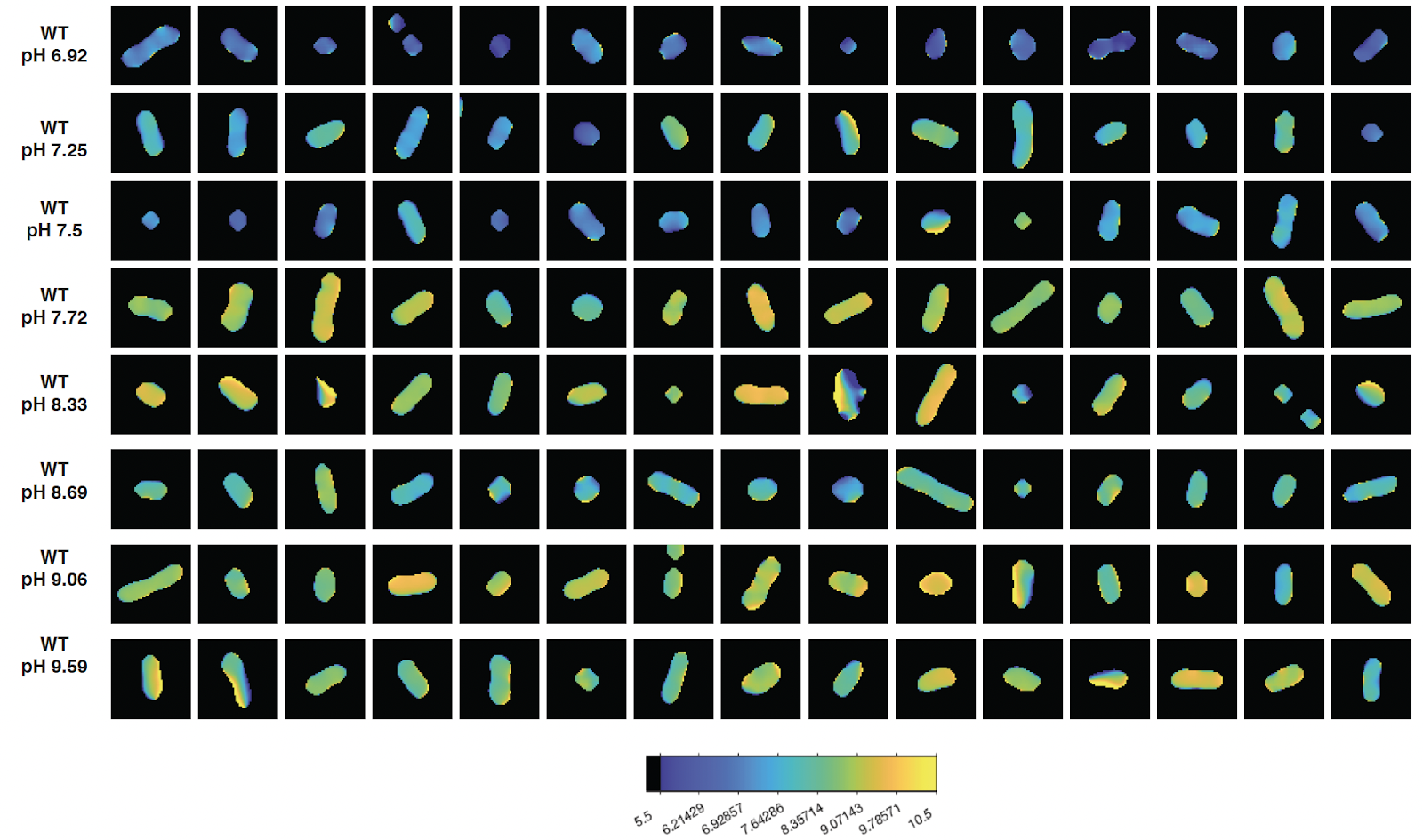

**Fig S8: Intracellular pH of WT cells across an environmental pH gradient.** Microscopic images of all replicate WT cells stained with BCECF in a range of pH adjusted buffers. Color of cells correspond to the cellular pH as denoted on the scale below microscopic images. Each image is cropped to 5.5 x 5.5 μm.

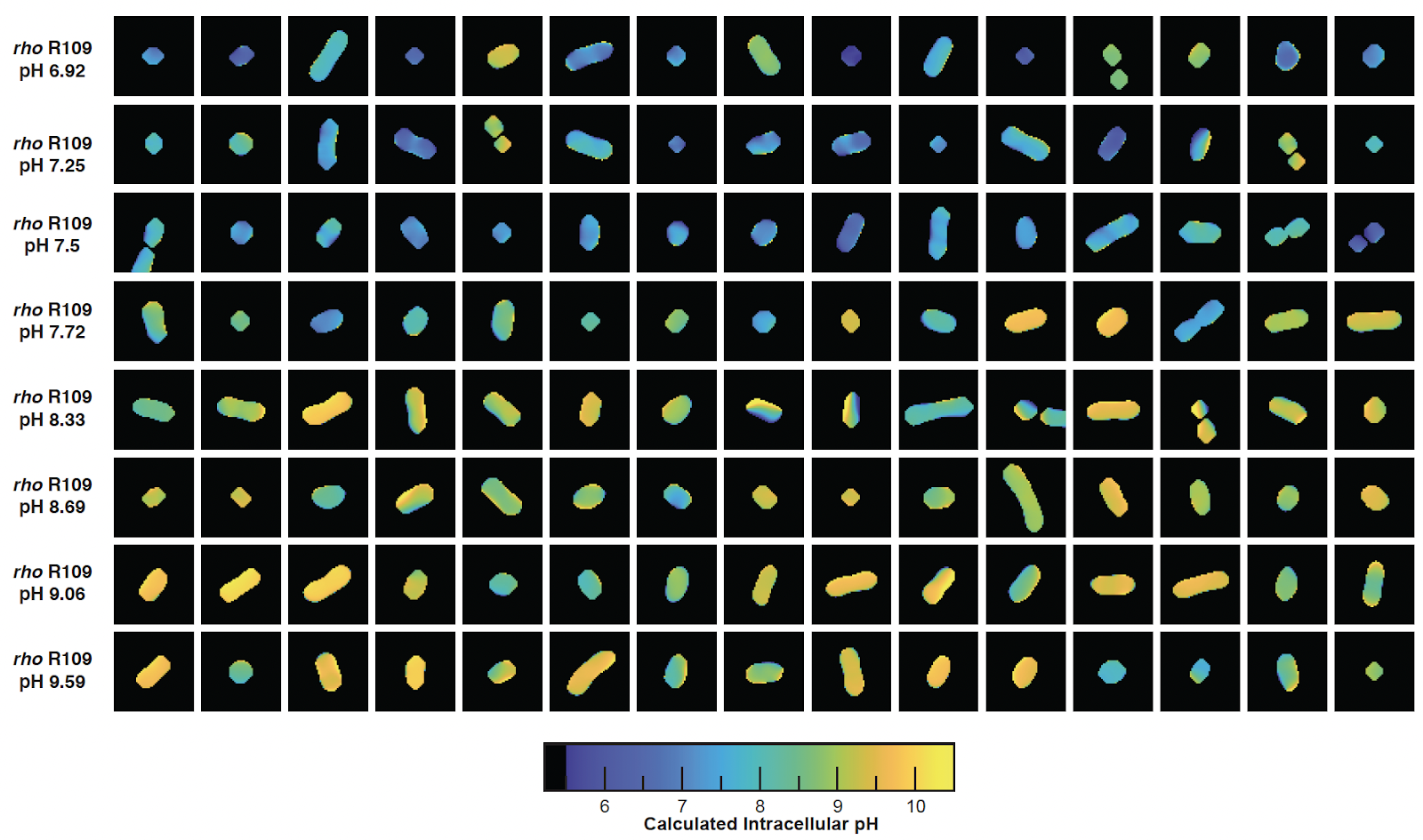

**Fig S9: Intracellular pH of *rho* R109H cells across an environmental pH gradient.** Microscopic images of all replicate *rho* R109H cells stained with BCECF in a range of pH adjusted buffers. Color of cells correspond to the cellular pH as denoted on the scale below microscopic images. Each image is cropped to 5.5 x 5.5 μm.

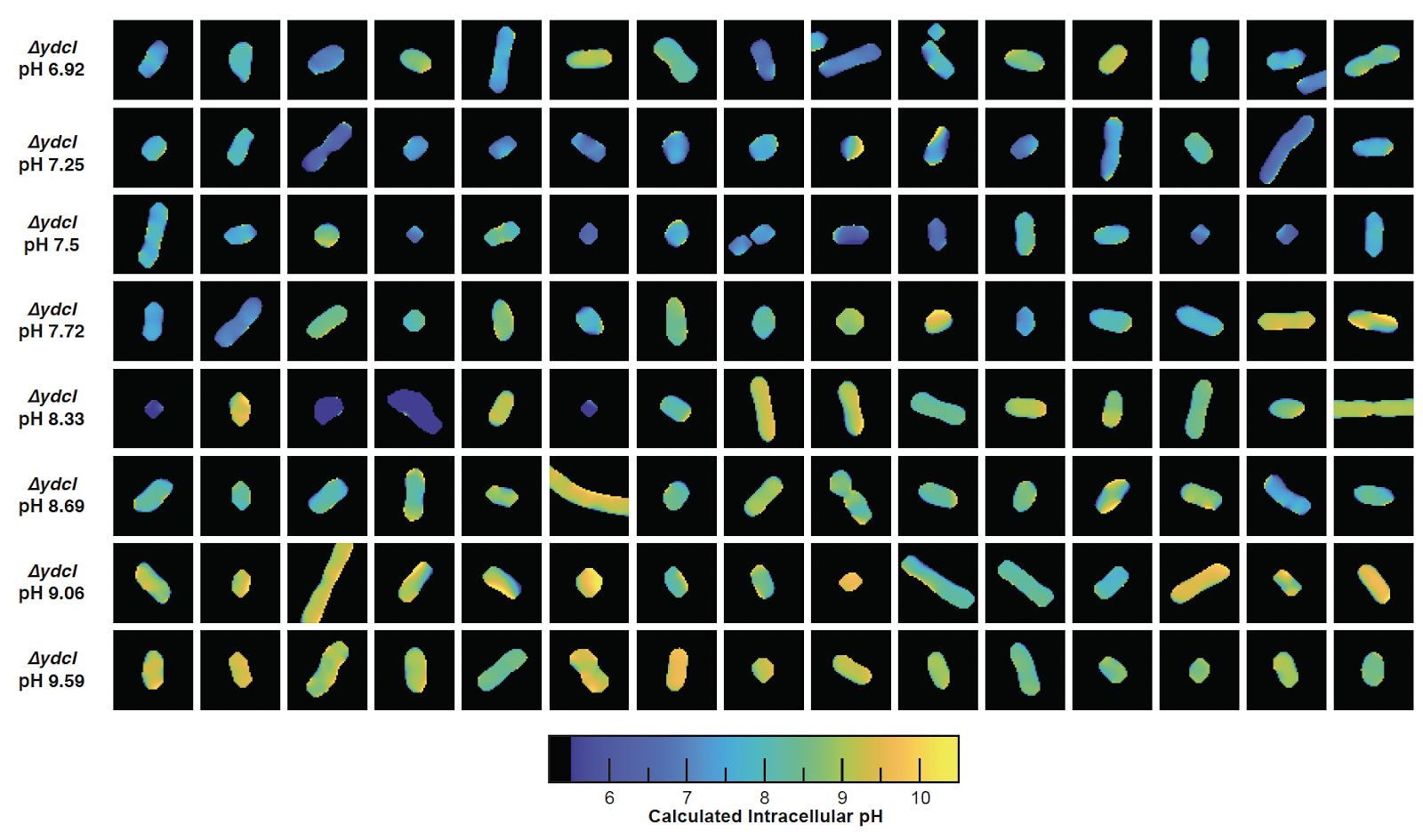

**Fig S10: Intracellular pH of** *Δ****ydcI* cells across an environmental pH gradient.** Microscopic images of all replicate *ΔydcI* cells stained with BCECF in a range of pH adjusted buffers. Color of cells correspond to the cellular pH as denoted on the scale below microscopic images. Each image is cropped to 5.5 x 5.5 μm.

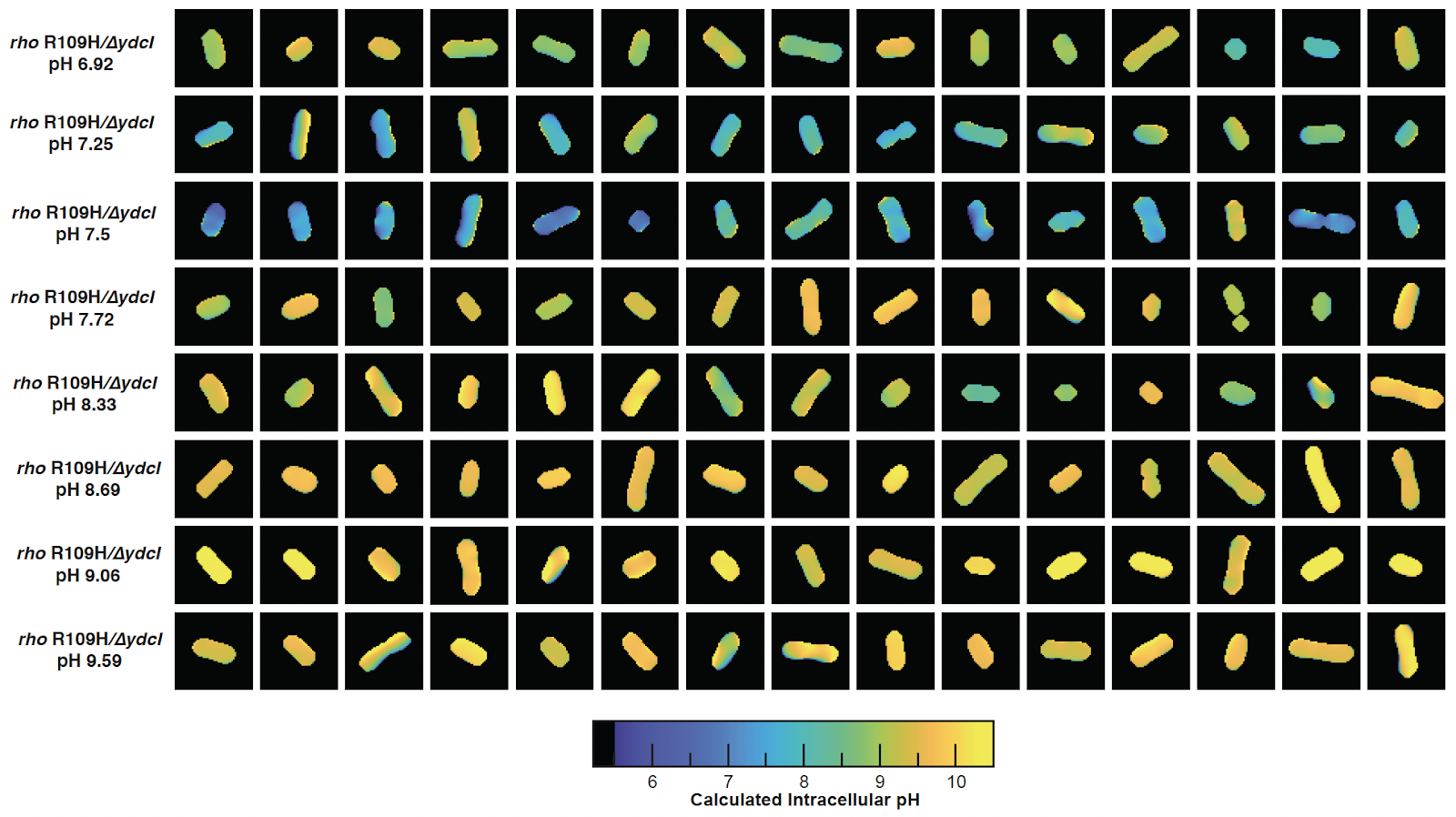

**Fig S11: Intracellular pH of *rho* R109H/***Δ****ydcI* cells across an environmental pH gradient.** Microscopic images of all replicate *rho* R109H/*ΔydcI* cells stained with BCECF in a range of pH adjusted buffers. Color of cells correspond to the cellular pH as denoted on the scale below microscopic images. Each image is cropped to 5.5 x 5.5 μm.

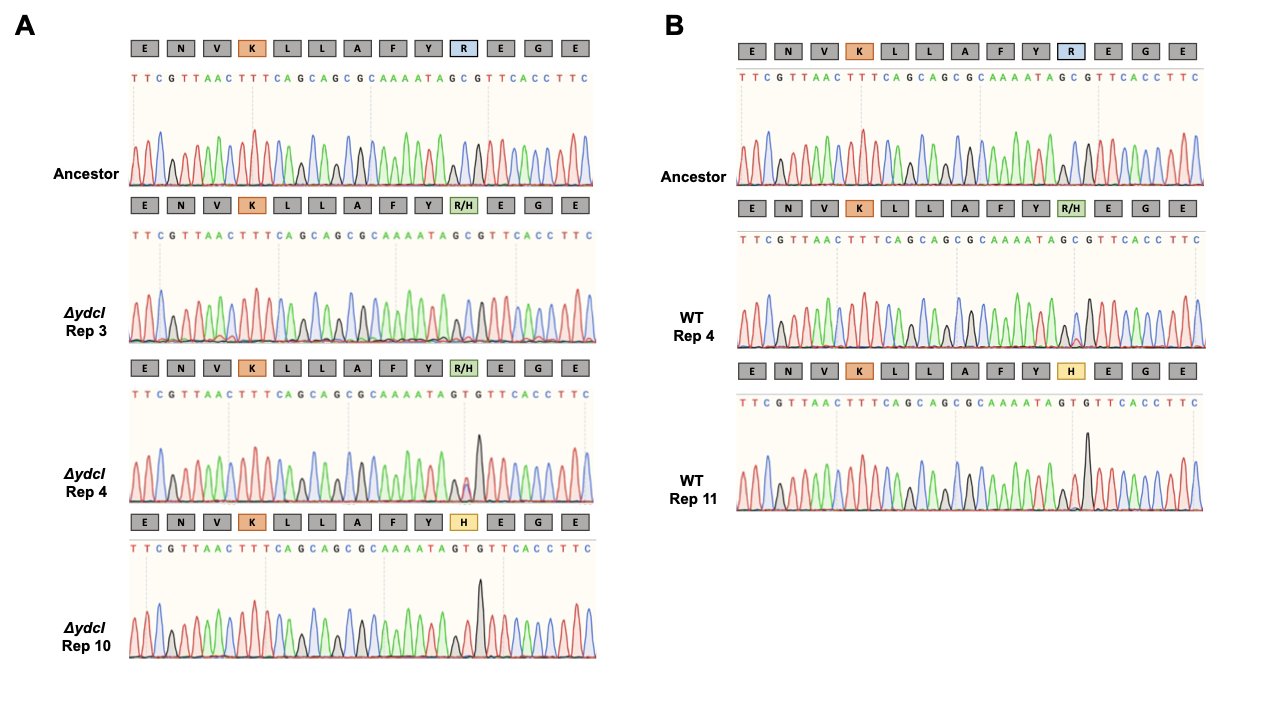

**Fig S12: Replaying one cycle of RLTS regime results in emergence of *rho* R109H mutation in initial *ΔydcI* and WT genetic background.** Portion of Sanger sequences containing the nucleotide sequence corresponding to the primary RNA binding site in Rho (Amino acid residues 106-118) with single letters above traces corresponding to the amino acid sequence. K105 residue is marked in orange, while R109 residue is marked in blue. The green R/H denotes a polymorphism at that position as indicated by the Sanger sequence. Yellow H denotes fixation of the R109H allele.

**Supplemental Tables**

| **Table S1.**  Relevant genotypes and phenotypes of populations adapted to 100-day feast/famine cycles | | | | |
| --- | --- | --- | --- | --- |
| **Population** | ***rho*** | ***ydcI*** | **AraBAD** | **MMR** |
| 501 | R109H | -3bp | AraBAD^+^ | MMR^+^ |
| 503 | K105E | -1bp | AraBAD^-^ | MMR^+^ |
| 504 | R109H | R10H | AraBAD^+^ | MMR^-^ |
| 506 | WT | V181A | AraBAD^-^ | MMR^-^ |
| 508 | WT | IS1 | AraBAD^-^ | MMR^+^ |
| 509 | WT | WT | AraBAD^+^ | MMR^-^ |
| 511 | R109H | D83G | AraBAD^-^ | MMR^-^ |
| 512 | WT | WT | AraBAD^+^ | MMR^+^ |
| 513 | WT | F74Y | AraBAD^+^ | MMR^+^ |
| 515 | WT | -1bp | AraBAD^-^ | MMR^+^ |
| 516 | G63S | R30C | AraBAD^+^ | MMR^-^ |
| 518 | R109H | +1bp | AraBAD^-^ | MMR^-^ |
| 520 | WT | L198Q | AraBAD^-^ | MMR^+^ |
| 521 | WT | WT | AraBAD^+^ | MMR^-^ |
| 523 | R109H | S91P | AraBAD^-^ | MMR^-^ |
| 524 | WT | -1bp | AraBAD^+^ | MMR^+^ |
